# Myeloid IRF5 is required for TLR7-driven inflammatory hemophagocyte differentiation and Macrophage Activation Syndrome

**DOI:** 10.64898/2026.08.18.745337

**Authors:** Natalie K. Thulin, Ailing Lu, Susana L. Orozco, Amanda Y.Y. Huang, LeAnn P. Nguyen, Gargi Mishra, Ram Savan, William L. Clapp, John P. Ray, Jessica A. Hamerman, Betsy J. Barnes

## Abstract

Inflammatory hemophagocytes (iHPCs) differentiate from Ly6C^HI^ monocytes, phagocytose red blood cells, and promote anemia and thrombocytopenia in a model of TLR7-driven macrophage activation syndrome (MAS), a life-threatening disorder. We identify IRF5 as a required and pharmacologically targetable driver of iHPC differentiation and MAS pathogenesis. Ly6C^HI^ monocyte-to-iHPC differentiation in TLR7.1 mice involved extensive transcriptome and chromatin remodeling. Gene programs gained in iHPCs were enriched for genes near accessible IRF5-binding regions that also carried NF-κB, ETS, and AP-1 motifs, suggesting collaborative regulation. IRF5 was required in vitro for TLR7-induced upregulation of a subset of these genes. Cell-intrinsic IRF5 expression was required for iHPC differentiation but dispensable for TLR7.1 Ly6C^HI^ monocytes and their transcriptional programs. Constitutive and myeloid-specific *Irf5* deletion, as well as a small-molecule IRF5 inhibitor, reduced iHPCs and ameliorated MAS and lupus features. IRF5 is therefore a broad regulator of iHPC differentiation and a tractable target in TLR7-driven inflammation.

**SUMMARY:** In TLR7-driven MAS, specialized monocyte-derived inflammatory hemophagocytes promote disease. Thulin, Lu et al. show that myeloid cell IRF5 is required for iHPC differentiation and MAS, and that IRF5 likely partners with other transcription factors, including NF-κB, to shape iHPCs programs.

## INTRODUCTION

Ly6C^HI^ inflammatory monocytes differentiate from myeloid progenitors in the bone marrow (BM) and, upon maturation, continuously enter circulation, where they respond to inflammatory and homeostatic cues. During homeostasis, Ly6C^HI^ monocytes support tissue repair and immune regulation^1^. During infection or injury, they initiate inflammatory responses, migrate to tissues, phagocytose pathogens, and differentiate into specialized effector populations^2^. When dysregulated, these responses can propagate sterile inflammation in autoimmune disease, cancer, and tissue injury^1,2^, underscoring the importance of understanding the mechanisms that regulate Ly6C^HI^ monocyte activation and differentiation. One such disease is macrophage activation syndrome (MAS), a life-threatening rheumatic condition characterized by fevers, cytopenias and the accumulation of activated macrophages that have phagocytosed red blood cells (RBCs) and leukocytes in blood-rich tissues^3^. MAS, and its counterpart secondary hemophagocytic lymphohistiocytosis, most commonly develops in patients with systemic juvenile idiopathic arthritis, systemic lupus erythematosus (SLE), hematopoietic malignancy, or viral infection^3^. Some MAS patients have mutations conferring heightened innate immune sensitivity or delayed resolution of adaptive immune responses^3–5^, and current models propose that MAS onset requires a combination of underlying inflammation, genetic predisposition, and an inflammatory triggering event, such as an infection or disease flare^5,6^. Identifying the signals that drive pathogenic macrophage development in MAS may inform effective, targeted therapeutics.

Endosomal TLRs (TLR3, 7, 8, 9) detect nucleic acids, and aberrant activation of these receptors by self-nucleic acids is a well-established driver of autoimmune diseases^7^. Among these, TLR7 is unequivocally linked to SLE pathogenesis, evidenced by gain-of-function mutations in *TLR7* or its signaling pathway components in patients^8–13^, as well as increased TLR7 engagement in mouse models of lupus-like disease^14,15^. Sustained endosomal TLR engagement in mice, through repetitive agonist injection, receptor overexpression, or loss of negative regulators, induces severe systemic inflammation and MAS-like disease, characterized by cytopenias and severe splenomegaly^15–20^. The TLR7.1 mouse, which overexpresses TLR7, develops spontaneous MAS-like disease secondary to lupus-like manifestations and serves as a valuable model for studying this pathology^14,15^. TLR signaling induces inflammatory cytokine production by myeloid cells and shapes myelopoiesis and monocyte fate. Chronic TLR activation promotes emergency myelopoiesis, characterized by inflammation-induced expansion of myeloid-biased progenitor cells, largely through cell-extrinsic effects on inflammatory cytokine production^21,22^. Cell-intrinsic TLR signaling in hematopoietic stem and progenitor cells can also promote macrophage differentiation in vitro^23–26^, and TLR-induced emergency myelopoiesis drives the generation of specialized inflammatory monocytes and macrophages^25,27^. In TLR7.1 mice, cell-intrinsic TLR7 signaling induces the differentiation of a unique Ly6C^HI^ monocyte-derived population not seen during homeostasis, termed inflammatory hemophagocytes (iHPCs), which accumulate in blood-rich tissues and phagocytose RBCs^15^. In TLR7.1 mice, the number of iHPCs containing internalized RBCs directly correlates with anemia, and ablation of Ly6C^HI^ monocytes, and consequent depletion of iHPCs, rescues mice from MAS-like disease.

Single-nucleotide polymorphisms in *IRF5,* encoding a transcription factor activated downstream of TLR7, are associated with increased risk of MAS, as well as several autoimmune diseases, including SLE, Sjögren’s disease, systemic sclerosis, and rheumatoid arthritis^28–33^. IRF5 is expressed broadly across immune populations and functions in a cell-type-specific manner, regulating B cell activation and plasmablast differentiation, as well as Th1 polarization^34–37^. In myeloid cells, IRF5 directly induces pro-inflammatory cytokines and type I interferons and regulates inflammatory macrophage differentiation^38–42^. *Irf5*-deficient mice accumulate significantly fewer iHPC-like cells after acute TLR7 agonist treatment; however, acute exposure does not recapitulate the complex differentiation of iHPCs or the full spectrum of MAS^15^. The specific contribution of myeloid IRF5 to MAS pathogenesis therefore remains undefined.

Here, we identified IRF5 as an essential driver of chronic TLR7-driven MAS-like disease and iHPC differentiation. Global and myeloid-specific genetic *Irf5* ablation, as well as therapeutic IRF5 inhibition in TLR7.1 mice, significantly reduced MAS severity and iHPC accumulation, establishing a requirement of myeloid cell IRF5 in iHPC development. Notably, our findings show that cell-intrinsic IRF5 expression is specifically required for the transition from TLR7.1 Ly6C^HI^ monocytes to iHPCs and define a distinct IRF5-regulated transcriptional program in iHPCs. Together, these findings establish myeloid IRF5 as a critical regulator of iHPC differentiation and a promising therapeutic target in MAS.

## RESULTS

### IRF5 drives core iHPC pathways but is dispensable for Ly6C^HI^ monocyte programs

To further understand the regulation of iHPC differentiation and function, we performed paired bulk RNA-seq and ATAC-seq on TLR7.1 monocyte populations and iHPCs sorted from chimeras reconstituted with TLR7.1 BM, which carries a ∼10-fold overexpression of *Tlr7* that drives chronic TLR7 signaling^14^ (Fig. 1 A, Fig. S1 A). iHPC RNA-seq and ATAC-seq profiles were distinct from those of Ly6C^HI^ monocytes from wild-type and TLR7.1 chimeras and other TLR7.1 monocyte populations by principal component analysis (Fig. 1 B). Differentially expressed genes (DEGs) and differentially accessible regions (DARs) between TLR7.1 Ly6C^HI^ monocytes and iHPCs revealed extensive epigenetic and transcriptional remodeling between the two populations (Fig. S2 A). We assigned each DAR to the nearest gene and found that DARs significantly correlated with expression of nearby DEGs (Fig. 1 C). To identify pathways and programs enriched in iHPCs and TLR7.1 Ly6C^HI^ monocytes, we performed over-representation analysis. Genes with increased expression and accessibility in iHPCs (Fig. 1 C, green, quadrant I) enriched for pathways characteristic of an inflammatory macrophage phenotype, including phagocytosis, cytokine production and inflammatory responses driven by NF-κB, and dynamic engagement with the immune environment (Fig. 1 D). In contrast, genes that had higher expression and accessibility in TLR7.1 Ly6C^HI^ monocytes (Fig. 1 C, blue, quadrant III) enriched for pathways associated with circulation, proliferative state, inflammatory innate immune responses, and interferon responses (Fig. 1 E).

**Figure 1:**
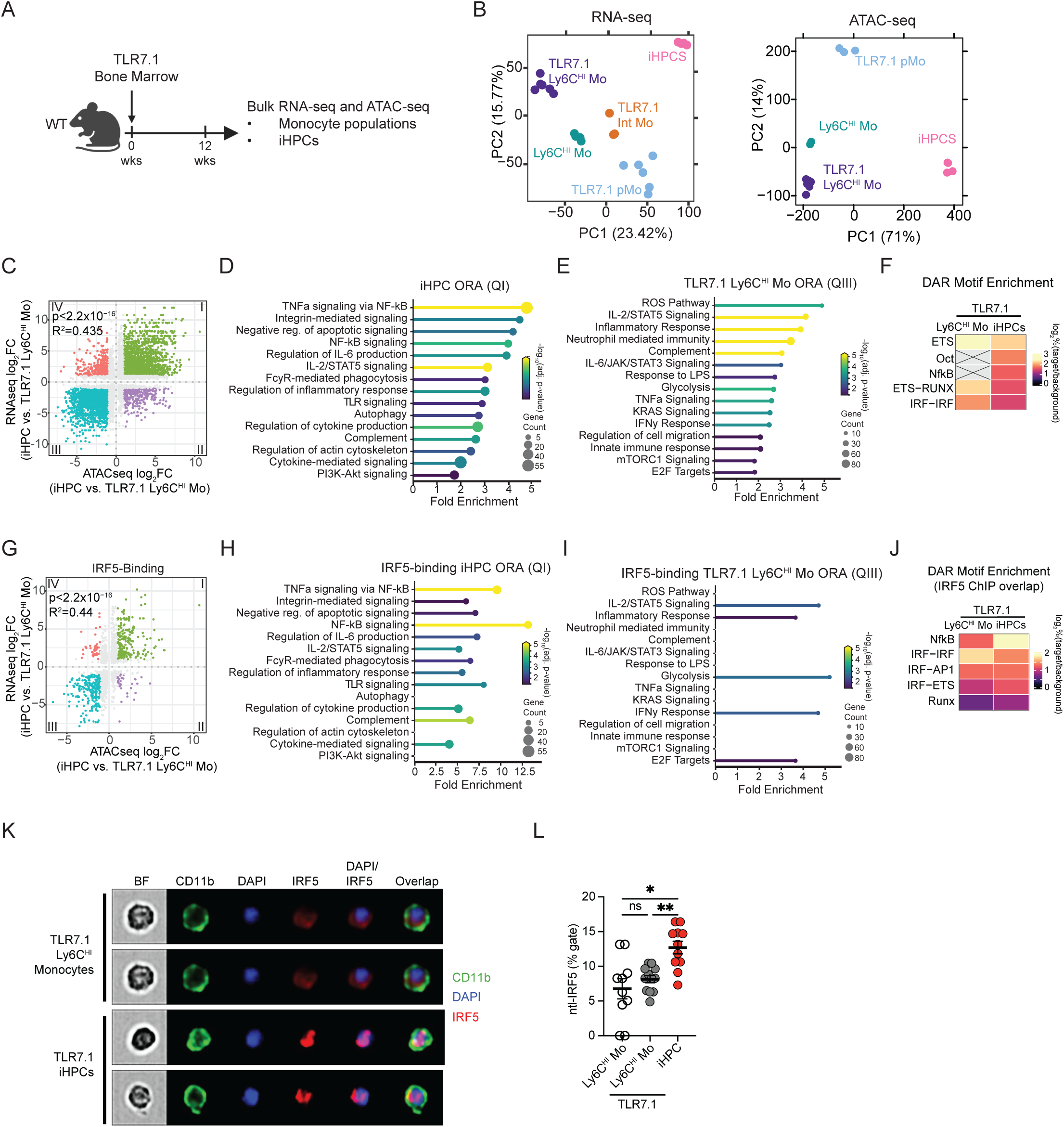
IRF5 drives an iHPC transcriptional and epigenetic program distinct from Ly6C^HI^ monocytes. (A-J) Paired bulk RNA sequencing and ATAC sequencing of splenic iHPCs (n=4) and monocyte populations (n=3-6) sorted from TLR7.1:TLR7.1 or WT:WT chimeras. (A) Experimental schematic. (B) Principal component analysis (PCA) plots of RNA-seq counts (left) and ATAC-seq counts (right) from all populations. (C) TLR7.1 Ly6C^HI^ monocyte and iHPC DAR and DEG correlation. Each dot represents a peak that corresponds to an accessible region near a gene that is differentially enriched (log_2_Fc ≥ 1, p < 0.05); R^2^ value and p value calculated on enriched peaks and genes (as shown in Fig. S2 A). The number of DARs mappings to DEGs in each quadrant: QI=2044, QII=339, QIII=1776, and QIV=369. (D) Over-Representation Analysis (ORA) of QI genes, upregulated with increased chromatin accessibility in iHPCs as compared to TLR7.1 Ly6C^HI^ monocytes (log_2_Fc ≥ 1, p < 0.05). (E) ORA of QIII genes, upregulated with increased chromatin accessibility in TLR7.1 Ly6C^HI^ monocytes as compared to iHPCs (log_2_Fc ≥ 1, p < 0.05). (F) Significantly enriched HOMER known motif families in the differentially accessible peaks between TLR7.1 Ly6C^HI^ monocytes and iHPCs, shown as frequency of the motif within open peaks (target) relative to frequency of the motif within all accessible peaks normalized for GC content (background). (G-J) IRF5 binding sites from IRF5 ChIP-seq data set^51^ were intersected with accessible regions in iHPCs and TLR7.1 Ly6C^HI^ monocytes. (G) Correlation of TLR7.1 Ly6C^HI^ monocyte and iHPC IRF5-binding DARs and DEGs. Each dot represents a peak that corresponds to an IRF5-binding accessible region near a gene that is differentially expressed (log_2_Fc ≥ 1, p<0.05); R^2^ value and p value calculated on enriched peaks and genes. The number of IRF5-binding DARs mapping to DEGs in each quadrant: QI=234, QII=47, QIII= 234, QIV=31. (H) ORA of IRF5-binding QI genes, upregulated iHPC genes with IRF5-binding accessible DARs (log_2_Fc ≥ 1, p<0.05). (I) ORA of IRF5-binding QIII genes, upregulated TLR7.1 Ly6C^HI^ monocytes genes with IRF5-binding accessible DARs (log_2_Fc ≥ 1, p<0.05). (J) HOMER known motif enrichment in IRF5 binding DAR peaks, shown as frequency of the motif within open peaks with published IRF5 binding (target) relative to frequency of the motif within non-IRF5-bound accessible peaks normalized for GC content (background). (K-L) IRF5 nuclear localization in Ly6C^HI^ monocytes and iHPCs from 9-week-old WT and TLR7.1 mice. Representative images (K) and quantification (L) of n=10-12 per group. *p<0.05, **p<0.01, Brown-Forsythe and Welch ANOVA test (L). Each symbol (L) represents individual mice; mean +/−SEM shown.

To define which transcription factors may be active in iHPCs and TLR7.1 Ly6C^HI^ monocytes, we used HOMER to screen open chromatin peaks from each cell type for enrichment of known transcription factor motifs. Because transcription factor families tend to share similar motifs, we clustered the HOMER identified transcription factors into 29 motif groups according to their motif similarity (Fig. S2 B). In more accessible DARs in both iHPCs and TLR7.1 Ly6C^HI^ monocytes, ETS family member motifs were the most enriched, consistent with their pioneering role in myeloid lineage commitment (Fig. 1 F). Strikingly, there was iHPC-specific enrichment of NF-κB motifs (Fig. 1 F), consistent with the NF-κB pathway enrichment observed in iHPC upregulated genes with more accessible DARs. Other iHPC-specific motif enrichment included Oct, NF-Y, and Nur77, all implicated in macrophage differentiation and inflammatory gene regulation (Fig. 1 F, Fig. S2 C). IRF-associated motifs, including single IRF and IRF-IRF homodimer, IRF-ETS composite, and IRF-AP1 composite motifs, were significantly enriched in both TLR7.1 Ly6C^HI^ monocytes and iHPCs; the latter two motifs are established drivers of myeloid differentiation and identity^43–45^ (Fig. 1 F, Fig. S2 C).

Given the enrichment of IRF motifs in iHPCs and the established role of IRF family members in myeloid lineages, we further explored IRF contribution to the iHPC-specific transcriptional and epigenetic program. IRF5 was of particular interest because it is activated downstream of TLR7 signaling and is associated with inflammatory macrophages^40–42,46–50^. Furthermore, we previously showed that IRF5 deficiency dampens the development of iHPC-like cells in vivo in response to acute TLR7 signaling ^15^. We therefore set out to directly assess the contribution of IRF5 to the iHPC transcriptional and epigenetic program. To directly link iHPC DARs to IRF5 occupancy we intersected our ATAC-seq data with a publicly available IRF5 ChIP-seq dataset from TLR9-stimulated dendritic cells^51^, finding 546 DARs identified in our dataset had known IRF5 binding sites. These TLR7.1 Ly6C^HI^ monocytes and iHPC DARs overlapping IRF5 ChIP-seq peaks were highly correlated with expression of nearby DEGs, with IRF5 binding in accessible chromatin associated with increased gene expression, consistent with IRF5 functioning as a transcriptional activator in these cells (Fig. 1 G). Genes associated with IRF5 binding that were upregulated and more accessible in iHPCs showed higher fold enrichment for many of the same pathways identified with the full dataset, suggesting that IRF5-binding DARs likely drive the iHPC-specific transcriptional program (Fig. 1 H). In contrast, in TLR7.1 Ly6C^HI^ monocytes, upregulated and accessible genes associated with IRF5 binding did not enrich for the TLR7.1 Ly6C^HI^ monocyte-specific pathways identified above (Fig. 1 I), suggesting that IRF5 is not an important regulator of TLR7.1 Ly6C^HI^ monocyte-specific transcriptional programs. HOMER transcription factor motif analysis of these DARs with known IRF5-binding regions in iHPCs and TLR7.1 Ly6C^HI^ monocytes identified in Fig. 1 G revealed enrichment over background for all IRF motif groups in both cell types as well as the highest enrichment for NF-κB motifs in iHPCs relative to TLR7.1 Ly6C^HI^ monocytes (Fig. 1 J, Fig. S2 D), consistent with the pathway analysis of iHPC IRF5 binding accessible regions strongly enriching for NF-κB pathways.

Having found that IRF5 occupancy selectively enriched for the iHPC-specific transcriptional program, we next asked whether IRF5 is preferentially activated in iHPCs in vivo. To assess the functional potential of IRF5 in iHPCs and Ly6C^HI^ monocytes, we quantified IRF5 activity in circulating myeloid cells of TLR7.1 mice. Because activated IRF5 translocates to the nucleus in response to TLR7 signaling^32,33,52^, we used imaging flow cytometry to assess nuclear IRF5 in wild-type (WT) and TLR7.1 mice. We detected equivalent low levels of nuclear-localized IRF5 in Ly6C^HI^ monocytes from WT and TLR7.1 mice, whereas iHPCs had significantly higher nuclear IRF5 than Ly6C^HI^ monocytes in both strains, indicating increased IRF5 activation (Fig. 1 K-L, Fig. S1 C). Together, these data suggest that IRF5 may be a key transcription factor in establishing and/or maintaining iHPC identity.

### TLR7-induced IRF5 signaling in Ly6C^HI^ monocytes is required for the induction of an iHPC gene program

To directly assess how IRF5 shapes the transition of Ly6C^HI^ monocytes to iHPCs, we used an in vitro culture system in which BM Ly6C^HI^ monocytes were treated with the TLR7 agonist R848. In this in vitro system, WT Ly6C^HI^ monocytes differentiate into iHPC-like cells, while *Irf5^−/−^*Ly6C^HI^ monocytes do not (Fig. S3 A-B). We conducted bulk RNA-sequencing of untreated and R848-treated WT and *Irf5^−/−^* Ly6C^HI^ monocytes at 6, 24, and 48 hours to capture early IRF5-dependent transcriptional changes (Fig. 2 A). Of the 2763 transcripts induced ≥ 2-fold by R848 in WT Ly6C^HI^ monocytes, 832 were significantly less induced in *Irf5^−/−^* Ly6C^HI^ monocytes (Fig. S3 C-D). To relate these genes to in vivo iHPC gene expression, we compared them to genes with increased expression and chromatin accessibility in iHPCs relative to Ly6C^HI^ monocytes from TLR7.1 mice (Fig. 1 C, green, quadrant I). This comparison revealed significant overlap, as of the 832 IRF5-dependent R848-induced genes in Ly6C^HI^ monocytes, 92 were also highly expressed in iHPCs in vivo (Fig. 2 B-C), demonstrating that iHPCs contain an IRF5-dependent gene signature.

**Figure 2:**
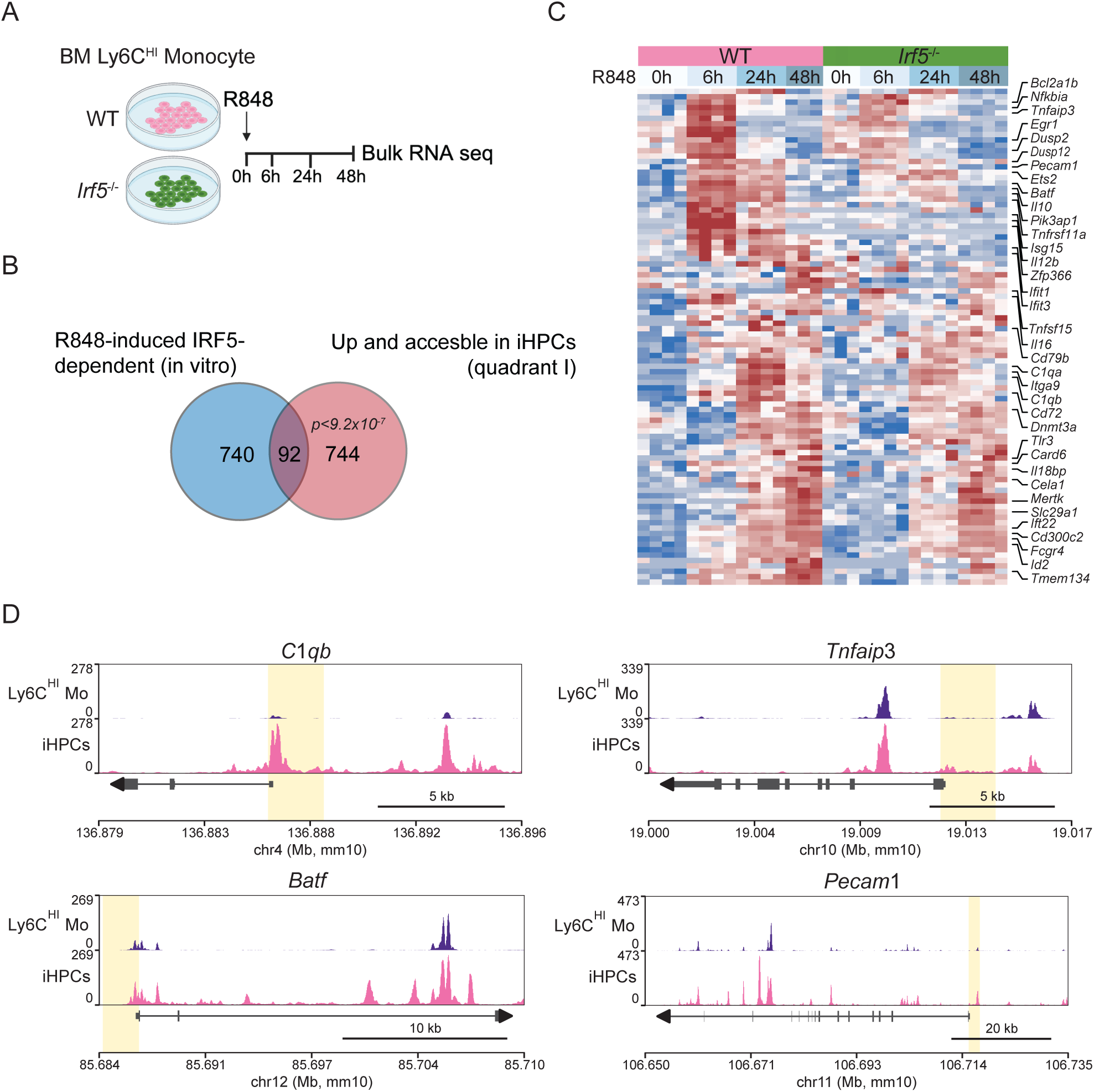
TLR7-induced IRF5 signaling in Ly6C^HI^ monocytes is required for the induction of an iHPC gene program. (A) WT and *Irf5^−/−^*BM Ly6C^HI^ monocytes were treated with M-CSF +/− R848 for 6, 24, or 48 hours and then analyzed by bulk RNA-seq (n=4 per group). (B) Gene overlap of IRF5-dependent R848-induced genes in BM Ly6C^HI^ monocytes in vitro and upregulated with increased chromatin accessibility in iHPCs as compared to TLR7.1 Ly6C^HI^ monocytes from Fig. 1 C, green dots, quadrant I. (C) Heatmap of the overlapping 92 genes from Fig. 2 B. (D) ATAC-seq tracks for select genes from Fig. 2 C. One-tailed exact hypergeometric test (B). Each column (C) represents individual mice.

Among these IRF5-dependent iHPC genes were the inflammatory cytokines *Il12b*, *Il16,* and *Tnfsf15*, the anti-inflammatory cytokine *Il10*, and *Il18bp*, which antagonizes IL-18 (Fig. 2 C, Fig S3 E). Both *Il12b* and Il10 are established targets of IRF5 in myeloid cells^33,38,39^. Further, elevated circulating IL-10 and IL18BP have both been reported in patients with MAS^53–55^. The genes *Nfkbia* and *Tnfaip3,* which encode negative regulators of inflammatory signaling, were also represented (Fig. 2 C-D, Fig S3 E). *Tnfaip3* is particularly notable for its strong association with autoimmune disease. We also found genes encoding proteins involved in phagocytosis, including *Fcgr4* and *Mertk,* a receptor critical for apoptotic cell clearance. IRF5-dependent iHPC genes included complement components *C1qa* and *C1qb*, which are upregulated in patients with MAS, and are elevated in *Irf5*-dependent inflammatory macrophages^40,56,57^ (Fig. 2 C-D, Fig S3 E). This gene set also included genes encoding receptors involved in immune cell migration and adhesion, including *Itga9* and *Pecam1*. *Pecam1,* which encodes CD31, is of particular interest, as iHPCs express high levels of CD31 in vivo and we use it as a marker of these cells^15^ (Fig. 2 C-D). Genes encoding transcription factors were also IRF5-dependent iHPC genes, notably *Ets2* and *Batf*, both associated with myeloid cell differentiation and inflammatory responses (Fig. 2 C-D, Fig. S3 E). Collectively, these data show that IRF5 contributes to TLR7-induced transcriptional programs in Ly6C^HI^ monocytes that are associated with the emergence of iHPC identity.

### IRF5 is required for iHPC differentiation and MAS-like disease in TLR7.1 mice

To investigate the in vivo requirement for IRF5 in iHPC differentiation and MAS, we generated TLR7.1 mice lacking *Irf5* (TLR7.1 *Irf5*^−/−^), heterozygous for *Irf5* (TLR7.1 *Irf5*^+/−^), or wild-type for *Irf5* (TLR7.1 *Irf5*^+/+^). We assessed iHPCs in the blood and spleen at 14-16 weeks of age^15^ (Fig. S1 C-D). Circulating iHPCs were abundant in TLR7.1 *Irf5*^+/+^ mice but were significantly reduced in TLR7.1 *Irf5*^−/−^and TLR7.1 *Irf5*^+/−^ mice (Fig. S4 A-B). This effect was even more pronounced in the spleen, where iHPCs composed 1-5% of all CD45^+^ splenic cells in TLR7.1 *Irf5*^+/+^ mice, but were nearly absent in TLR7.1 *Irf5*^−/−^ and TLR7.1 *Irf5*^+/−^ mice (Fig. 3 A-B). Ly6C^HI^ monocytes were expanded in TLR7.1 mice relative to WT mice^15,26^, and *Irf5* deficiency reduced this expansion (Fig. 3 B, Fig. S4 B). We also assessed if IRF5 regulates iHPC phagocytosis of RBCs, a key function of these cells assessed by intracellular staining for the RBC marker TER-119^15^ (Fig. 3 C, Fig. S1 C-D). iHPC RBC phagocytosis was associated with *Irf5* gene dosage in both blood and spleen, with little hemophagocytosis when TLR7.1 mice lacked 1 or 2 copies of IRF5 (Fig. 3 C-D, S4 C). Blood, but not splenic, Ly6C^HI^ monocytes had high levels of RBC phagocytosis in TLR7.1 mice, which was reduced to WT levels with loss of one or two copies of *Irf5* (Fig. 3 D, S4 C). Together, these data demonstrate that IRF5 is essential for iHPC differentiation and/or accumulation as well as function in TLR7.1 mice, with even *Irf5* haploinsufficiency resulting in a dramatic reduction in iHPC number and hemophagocytosis.

**Figure 3:**
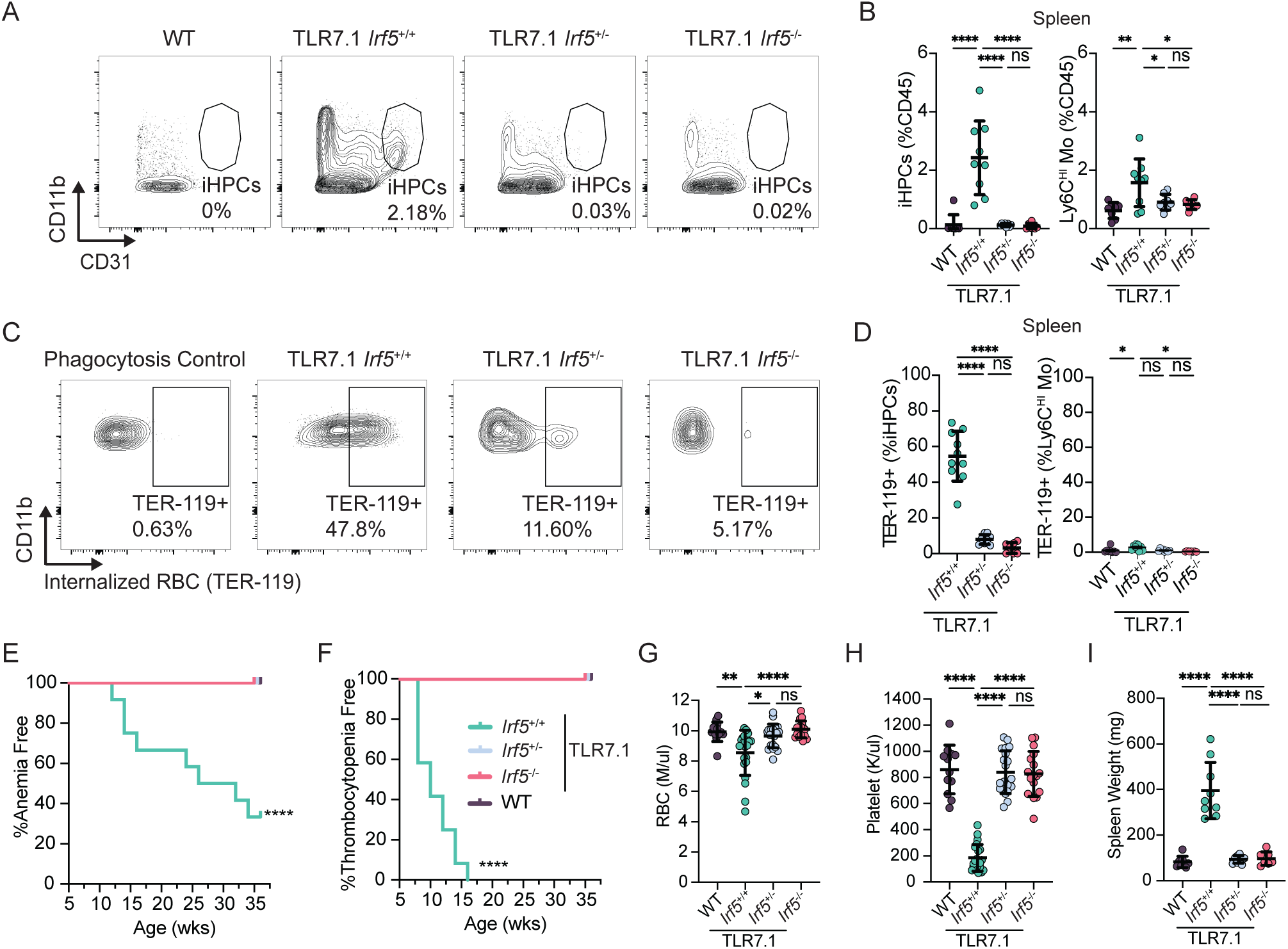
IRF5 is required for iHPC differentiation and function and MAS in vivo. (A-D) WT, TLR7.1 *Irf5*^+/+^, TLR7.1 *Irf5*^+/−^, and TLR7.1 *Irf5*^−/−^mice were assessed for iHPC development and RBC phagocytosis by flow cytometry in the spleen at 14-16 weeks of age (n=8-10 per group). (A) Representative gating of splenic iHPCs as a percent of CD45^+^ cells after gating on live CD45^+^Ly6G^−^SSC-A^LO^F4/80^−^ cells (full gating shown in Fig. S1 D). (B) Splenic iHPCs and Ly6C^HI^ monocytes as percent of CD45^+^ cells. (C) Representative gating of splenic iHPC RBC phagocytosis measured by intracellular TER-119 staining; a purified TER-119 block was used as an RBC phagocytosis staining control and to set the gate for internalized RBCs (TER-119^+^ cells). Splenic iHPCs were gated as in (A). (D) Percentage of splenic iHPCs and Ly6C^HI^ monocytes with intracellular RBCs. (E-I) WT, TLR7.1 *Irf5*^+/+^, TLR7.1 *Irf5*^+/−^, and TLR7.1 *Irf5*^−/−^ mice were assessed for anemia and thrombocytopenia every 2 weeks from 6-36 weeks of age (n=12-14 per group). Thrombocytopenia, platelets <400 K/μl; anemia, RBC <7 M/μl. Kaplan-Meier curves showing the percentage of mice that did not develop anemia (E) or thrombocytopenia (F). (G-I) Mice were assessed at 14-16 weeks of age for RBC count (G), platelet count (H), and spleen weight (I), (n=12-22 per group).*p< 0.05, **p<0.01, ****p<0.0001, Kruskal-Wallis test (B, D, G-I), Log-rank (Manel-cox) test (E-F); each symbol represents an individual mouse; mean +/− SEM shown.

We next asked if *Irf5* deficiency protects against the MAS-like disease that develops in TLR7.1 mice. In a longitudinal cohort, 60% of TLR7.1 *Irf5*^+/+^ mice developed moderate to severe anemia (RBCs < 7 M/μl) by 36 weeks of age, and all had developed severe thrombocytopenia (platelets < 400 K/μl) by 16 weeks. Fatal MAS, characterized by severe anemia (RBC < 5M/μl), emerged in a subset of TLR7.1 *Irf5*^+/+^ mice beginning at 18 weeks of age, however the loss of one or both *Irf5* alleles completely protected TLR7.1 mice from both anemia and thrombocytopenia (Fig. 3 E-H) as well as the severe splenomegaly characteristic of TLR7.1 mice^14,15^ (Fig. 3 I). We also assessed the impact of *Irf5* loss on the underlying lupus-like disease in the TLR7.1 mice. Loss of one or two copies of IRF5 fully protected TLR7.1 mice from lupus-like disease, with levels of anti-RNA autoantibodies and glomerulonephritis reduced to that similar to WT mice as were accumulations of activated T cells and alterations in B cell populations in the spleen (Fig. S4 D-I), similar to the required role for IRF5 in other murine lupus models ^14,35,58–60^. Together, these data demonstrate that loss of a single *Irf5* allele is sufficient to protect TLR7.1 mice from both MAS-like and lupus-like disease.

### Cell-intrinsic IRF5 expression is required for iHPC differentiation

To disentangle whether the lack of iHPCs in TLR7.1 *Irf5*^−/−^ mice is due to a cell-intrinsic requirement for IRF5 in iHPC differentiation or if it is due to the lack of lupus-like disease and overall inflammation in these mice, we generated mixed BM chimeras in which B6.CD45.1 recipient mice were reconstituted with an equal mixture of TLR7.1 (CD45.1.2) and TLR7.1 *Irf5*^−/−^ (CD45.2) donor BM distinguished by congenic markers. In parallel, control chimeras were generated using TLR7.1 BM from two congenically distinct donors (Fig. 4 A). In TLR7.1:TLR7.1 *Irf5*^−/−^ chimeras, splenic iHPCs were almost exclusively derived from *Irf5*-sufficient donor cells showing that *Irf5*-deficient iHPCs had a strong disadvantage. Control chimeras had similar iHPC reconstitution from each congenically distinct TLR7.1 donor. In contrast to iHPCs, splenic Ly6C^HI^ monocytes reconstituted equally in experimental and control chimeras (Fig. 4 B-D). TLR7.1 *Irf5*^−/−^ iHPCs were also at significant disadvantage in blood (Fig. 4 E). These data demonstrate a cell-intrinsic requirement for IRF5 in iHPC differentiation. In contrast, IRF5 is dispensable for TLR7.1 Ly6C^HI^ monocyte reconstitution, indicating that the IRF5-dependent monocytosis (Fig. 3B, Fig. S4 B) observed in TLR7.1 mice is caused by IRF5-mediated cell extrinsic effects likely on global inflammation.

**Figure 4:**
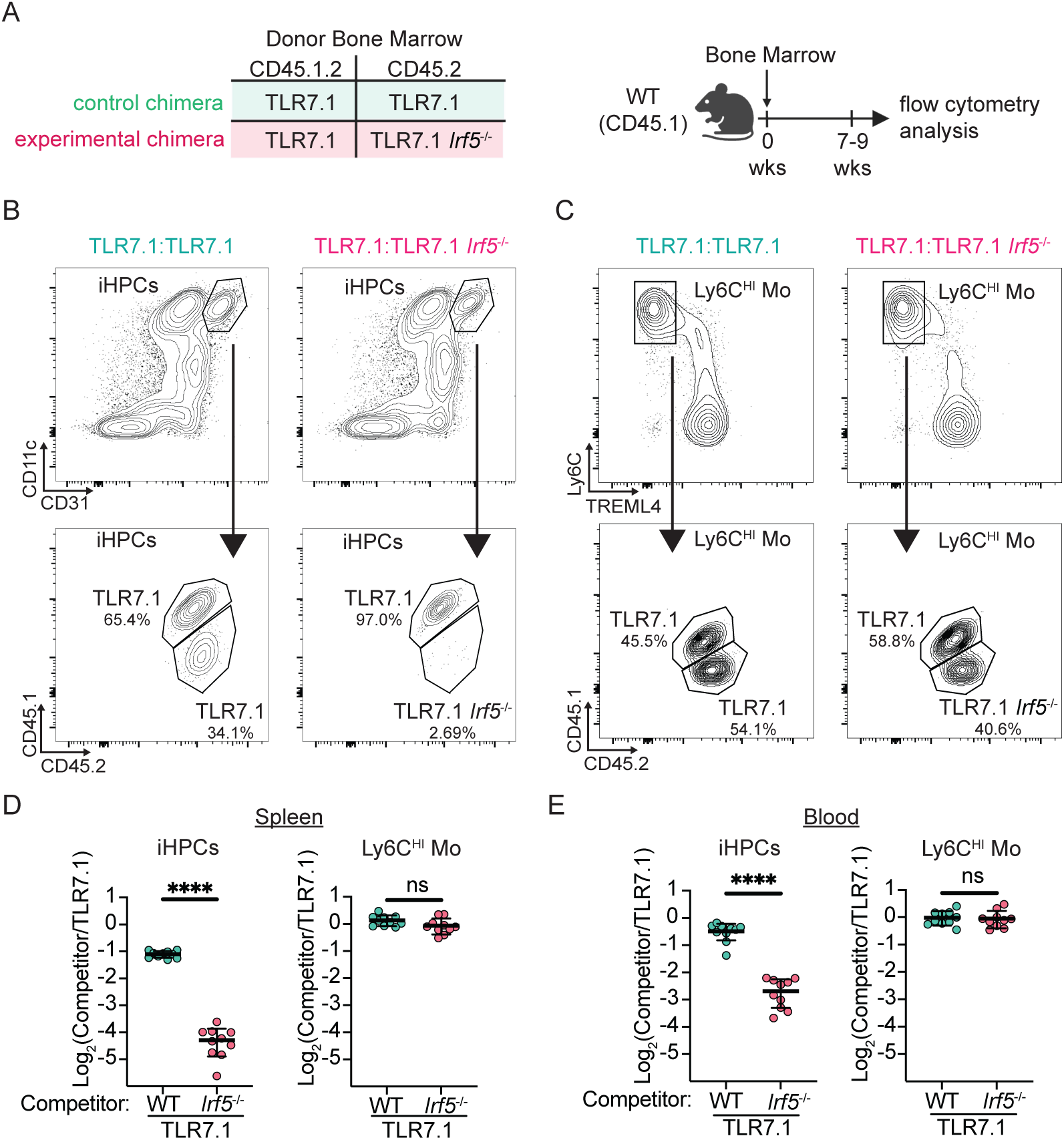
Cell-intrinsic IRF5 expression is required for iHPC differentiation. (A) Mixed BM chimeras were generated by reconstituting B6.CD45.1 donor mice with a 1:1 ratio of TLR7.1 *Irf5*^−/−^ (CD45.2) and TLR7.1 (CD45.1.2) BM; control chimeras were reconstituted with a 1:1 ratio of TLR7.1 (CD45.2) and TLR7.1 (CD45.1.2) BM. (B-C) Representative flow cytometry identifying CD45.2 and CD45.1.2 splenic (B) iHPCs and (C) Ly6C^HI^ monocytes in chimeras reconstituted for 7-9 weeks. (D-E) Ratio of CD45.2 (TLR7.1 *Irf5*^−/−^ or TLR7.1) to CD45.1.2 (TLR7.1) cells in chimeras for (D) splenic and (E) blood iHPCs and Ly6C^HI^ monocytes. Data are representative of three experiments; each point is an individual mouse, mean +/−SEM shown (n=10 per group). ****p<0.0001, Student’s unpaired t-test.

When we examined the cell-intrinsic contribution of IRF5 to other immune populations we found that other splenic myeloid cell populations including neutrophils and red pulp macrophages reconstituted similar to control chimeras (Fig. S5 A). T cell reconstitution was mildly promoted in the absence of *Irf5* expression (Fig. S5 B). However, several B cell populations showed a cell-intrinsic requirement for IRF5, including germinal center B cells and immature/transitional B cells (Fig. S5 C). Together, these data show that IRF5 does not globally regulate immune cell differentiation during chronic TLR7 signaling. Instead, cell-intrinsic IRF5 is selectively required for the Ly6C^HI^ monocyte to iHPC transition and also promotes the differentiation of specific B cell populations.

Because TLR7.1 *Irf5*^−/−^ Ly6C^HI^ monocytes fail to differentiate into iHPCs in mixed BM chimeras, we examined whether IRF5 poises TLR7.1 Ly6C^HI^ monocytes for iHPC differentiation. Paired bulk RNA-seq and ATAC-seq was performed on TLR7.1 Ly6C^HI^ monocytes and TLR7.1 *Irf5*^−/−^ Ly6C^HI^ monocytes isolated from TLR7.1:TLR7.1 *Irf5*^−/−^ mixed BM chimeras, ensuring the cells experienced the same inflammatory milieu. RNA-seq and ATAC-seq profiles from TLR7.1 Ly6C^HI^ monocytes and TLR7.1 *Irf5*^−/−^ Ly6C^HI^ monocytes were virtually identical, with no detectable DEGs and very few DARs between the two populations (Fig. S5 D). These findings further suggest that IRF5 is dispensable for maintaining the transcriptional and chromatin state of TLR7.1 Ly6C^HI^ monocytes but is instead required for their differentiation into iHPCs.

### Myeloid IRF5 expression is required for iHPC development and MAS

While IRF5 acts across multiple immune compartments, including in B cells ^59^, to promote lupus-like disease, the specific role of myeloid cell IRF5 in MAS has not been investigated. We therefore examined the role of myeloid-expressed IRF5 in iHPC differentiation and MAS-like disease, independent of any effect of IRF5 on B cells and other lymphocytes. Because female TLR7.1 mice developed anemia at a higher rate than male TLR7.1 mice in our previous experiments (anemic: female, 5/5; male, 3/7), we used female mice for these experiments. We crossed TLR7.1 mice with *Lyz2*^cre/cre^*Irf5*^Fl/Fl^ mice (denoted TLR7.1 *Irf5*^ΔM^) to delete *Irf5* from key myeloid populations, including Ly6C^HI^ monocytes and their progeny, macrophages, neutrophils, and some dendritic cell populations^61^. To control for the loss of lysozyme M that occurs with the homozygous *Lyz2^Cre^* allele, we generated TLR7.1 *Lyz2*^cre/cre^*Irf5*^WT/WT^ (denoted TLR7.1 *Irf5*^WT^) mice as controls. By qRT-PCR analysis of *Irf5* mRNA in these mice, we found significant, but incomplete, reduction of *Irf5* in Ly6C^HI^ monocytes, a large reduction in neutrophils and no change in total lymphocytes or B cells (Fig. S6 A-B). Strikingly, at 14-18 weeks of age, we found a dramatic reduction in splenic iHPCs in TLR7.1 *Irf5*^ΔM^ mice as compared with TLR7.1 *Irf5*^WT^ controls (Fig. 5 A-B). The few iHPCs that remained in TLR7.1 *Irf5*^ΔM^ mice were able to phagocytose RBCs and did not show any reduction in *Irf5* mRNA, suggesting they differentiated from the pool of Ly6C^HI^ monocytes without *Irf5* deletion (Fig. S6 A, C). Splenic Ly6C^HI^ monocytes were also significantly reduced in TLR7.1 *Irf5*^ΔM^ mice, comparable to WT control levels (Fig. 5 B). These findings show that IRF5 acts within the myeloid compartment to promote iHPC differentiation.

**Figure 5:**
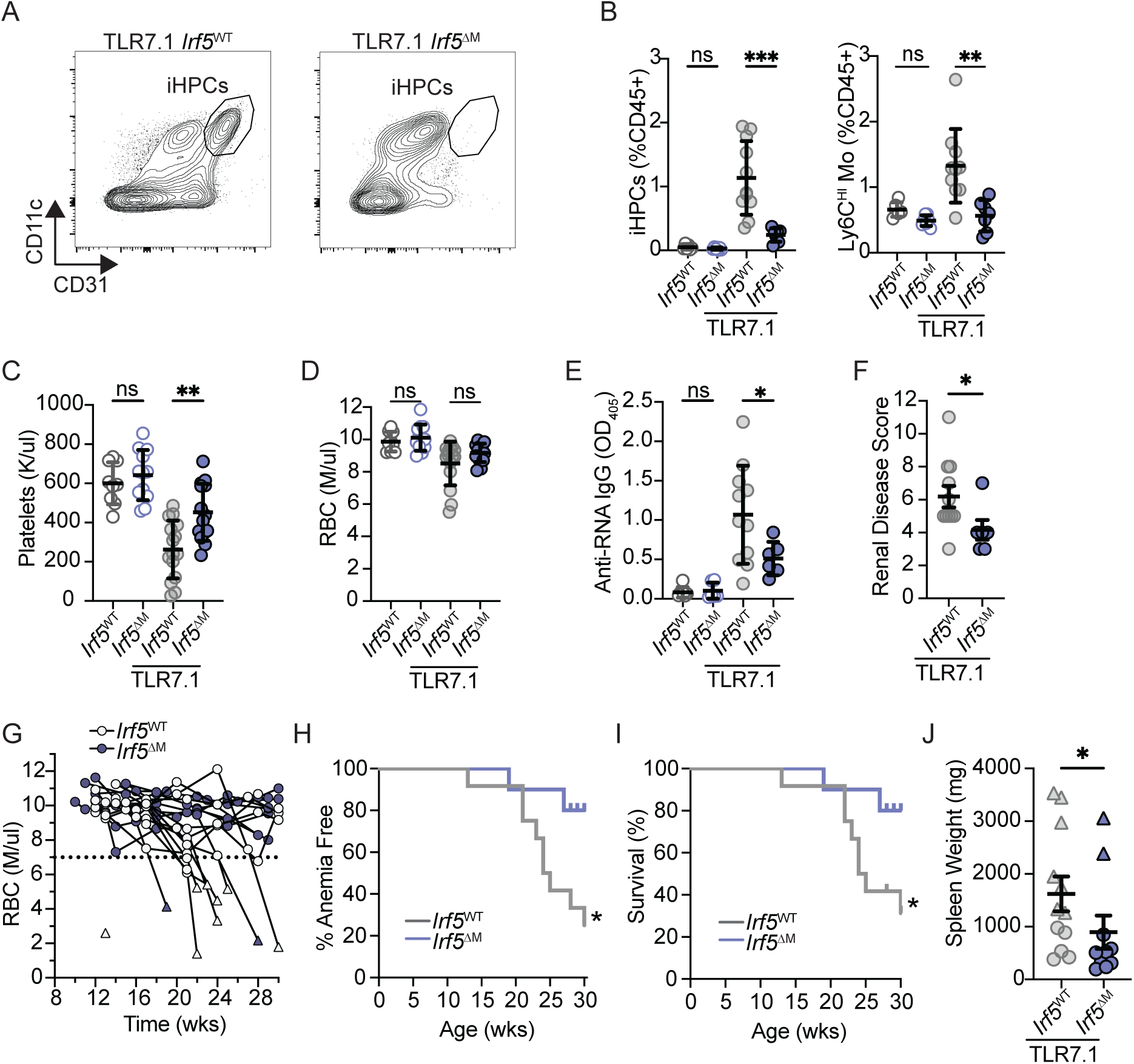
Myeloid *Irf5* expression is required for iHPC differentiation and MAS. (A-F) Female TLR7.1 *Irf5*^ΔM^ (*Lyz2*^CRE/CRE^ *Irf5*^FL/FL^) and TLR7.1 *Irf5*^WT^ (*Lyz2*^CRE/CRE^*Irf5*^WT/WT^) mice were assessed for iHPCs, MAS-like and lupus-like disease at 14-18 weeks of age (n=6-11 per group). (A) Representative flow cytometry of splenic iHPCs of gated CD45^+^F4/80^−^Ly6G^−^SiglecF^−^CD11b^+^ cells (gated as in Fig. S1 A, excluding CD115). (B) Splenic iHPCs and Ly6C^HI^ monocytes as percentage of CD45^+^ cells. (C) Platelet counts, (D) RBC counts, (E) serum anti-RNA IgG, and (F) renal disease score. (G-J) TLR7.1 *Irf5*^ΔM^ and TLR7.1 *Irf5*^WT^ mice were assessed every 3-4 weeks from 10-30 weeks of age (n=10-12 per group). (G) RBC counts; the dotted line denotes the cutoff for anemia; triangle represents terminal point. (H, I) Kaplan-Meier curves showing the percentage of mice that did not develop anemia (H) or succumb to disease (I) over the course of the experiment. (J) Spleen weights at 30 weeks or time of euthanasia; triangles represent mice that were euthanized due to MAS prior to 30 weeks and circles represent mice euthanized at 30 weeks of age. *p<0.05, **p<0.01, ***p<0.001, Brown-Forsythe and Welch ANOVA test (B, C), Kruskal-Wallis test (C, E), Welch’s t-test (F, J), Log-rank (Manel-cox) test (H, I). (B-G, J) each symbol is an individual mouse; mean +/−SEM shown.

We then assessed measures of disease. At 14-18 weeks of age, TLR7.1 *Irf5*^ΔM^ mice had significantly higher platelet counts than TLR7.1 *Irf5*^WT^ mice, showing that the loss of myeloid *Irf5* reduced or delayed development of thrombocytopenia (Fig. 5 C). There was little anemia yet at this time point (Fig. 5 D). Unexpectedly, TLR7.1 *Irf5*^ΔM^ mice had reduced serum anti-RNA autoantibodies and reduced early glomerulonephritis, even though the overall B cell and T cell compartments were unchanged by myeloid *Irf5* deficiency (Fig. 5 E-F, Fig. S6 D-F). Because the few mice that developed anemia and severe splenomegaly at this early time were all from the TLR7.1 *Irf5*^WT^ group, we extended disease assessment in TLR7.1 *Irf5*^WT^ and TLR7.1 *Irf5*^ΔM^ mice to 28-30 weeks of age. We found that TLR7.1 *Irf5*^ΔM^ mice were significantly protected from anemia, had significantly improved survival, and reduced splenomegaly compared to TLR7.1 *Irf5*^WT^ mice (Fig. 5 G-J). Together these data support a requirement for myeloid IRF5 in iHPC development in TLR7.1 mice and show that myeloid *Irf5* deletion protects against MAS-like disease.

### IRF5 inhibition blunts iHPC differentiation and ameliorates MAS

Because our genetic models showed that IRF5 is critical for iHPC and MAS development, we explored whether targeting IRF5 would be efficacious therapeutically. We leveraged a cell-permeable, non-toxic IRF5 inhibitor, N5-1, which selectively binds the inactive IRF5 monomer and prevents nuclear translocation without blocking other aspects of TLR7 signaling^62^. The N5-1 inhibitor has demonstrated efficacy in SLE, ameliorating multiple disease pathologies including antinuclear antibodies, circulating plasma cells, glomerulonephritis, and survival^62^. We first tested whether N5-1 treatment before mice show features of MAS could reduce iHPC development and prevent or delay MAS in TLR7.1 mice. Inhibitor treatment started at 5 weeks of age, with 10 doses over 6 weeks (Fig. S7 A). In inhibitor treated mice compared to vehicle treated controls, we found reduced circulating iHPCs at 13 and 15 weeks of age, when iHPCs typically peak (Fig. S7 B). In a larger cohort assessed at 17 weeks of age, 6 weeks after cessation of treatment, N5-1 treated mice had a significantly reduced circulating and splenic iHPCs, as well as iHPCs with internalized RBCs, compared to vehicle treated mice (Fig. S7 C-E). At this 17 week timepoint, splenomegaly showed a trending reduction, however anemia was not prevented (Fig S7 F-G). Crucially, early treatment with the IRF5 inhibitor N5-1 significantly reduced iHPC expansion (Fig. S7 B-C, E) and reduced mortality in TLR7.1 mice out to 40 weeks of age (Fig. S7 H), collectively demonstrating its protective effect against key MAS features.

To assess the therapeutic potential of IRF5 inhibition after disease onset, we administered 10 doses of the N5-1 inhibitor to TLR7.1 mice over 6 weeks, beginning at 13 weeks of age. This age of treatment initiation was chosen because mice typically exhibited thrombocytopenia but not yet anemia, indicating early MAS-like disease (Fig. 3 E-F, Fig. 6 A). This treatment regimen targets IRF5 activity in the weeks leading up to and during typical disease development, characterized by peak iHPC numbers and onset of anemia (Fig. 3 F). N5-1 treated mice had significantly reduced blood iHPCs at 16 and 18 weeks of age compared to vehicle treated mice (Fig. 6 B, C). At 18 weeks of age, we also found reduced blood iHPCs with internalized RBCs in N5-1 treated TLR7.1 mice (Fig. 6 D). To determine if the efficacy of this treatment extended to splenic iHPCs and MAS-associated tissue pathology, we assessed a separate cohort of mice at 17 weeks of age, 4 weeks into this treatment protocol. At this timepoint, both the total number of splenic iHPCs and splenomegaly were significantly reduced in the N5-1 treated mice (Fig. 6 E-F). Through 18 weeks of age, fewer N5-1 treated mice developed moderate to severe anemia compared to vehicle treated controls (Fig. 6 G). We also examined whether therapeutic treatment provided long term protection to TLR7.1 mice. IRF5 inhibitor treated mice had improved survival compared to vehicle-treated TLR7.1 mice through 40 weeks of age, even though the 6-week inhibitor treatment ended at 19 weeks (Fig. 6 H). Together, these data show that both preventative and therapeutic IRF5 inhibitor treatment reduce iHPC numbers and RBC phagocytosis, ameliorate aspects of MAS, and protect against mortality in TLR7.1 mice, with the effect on splenomegaly and anemia more pronounced with therapeutic inhibitor treatment, which spans a later time during disease. Given the robust reduction in circulating and total splenic iHPCs, as well as in iHPC RBC phagocytosis, we hypothesize that reducing IRF5 activity in iHPCs contributes to the protection observed in IRF5 inhibitor treated mice.

**Figure 6:**
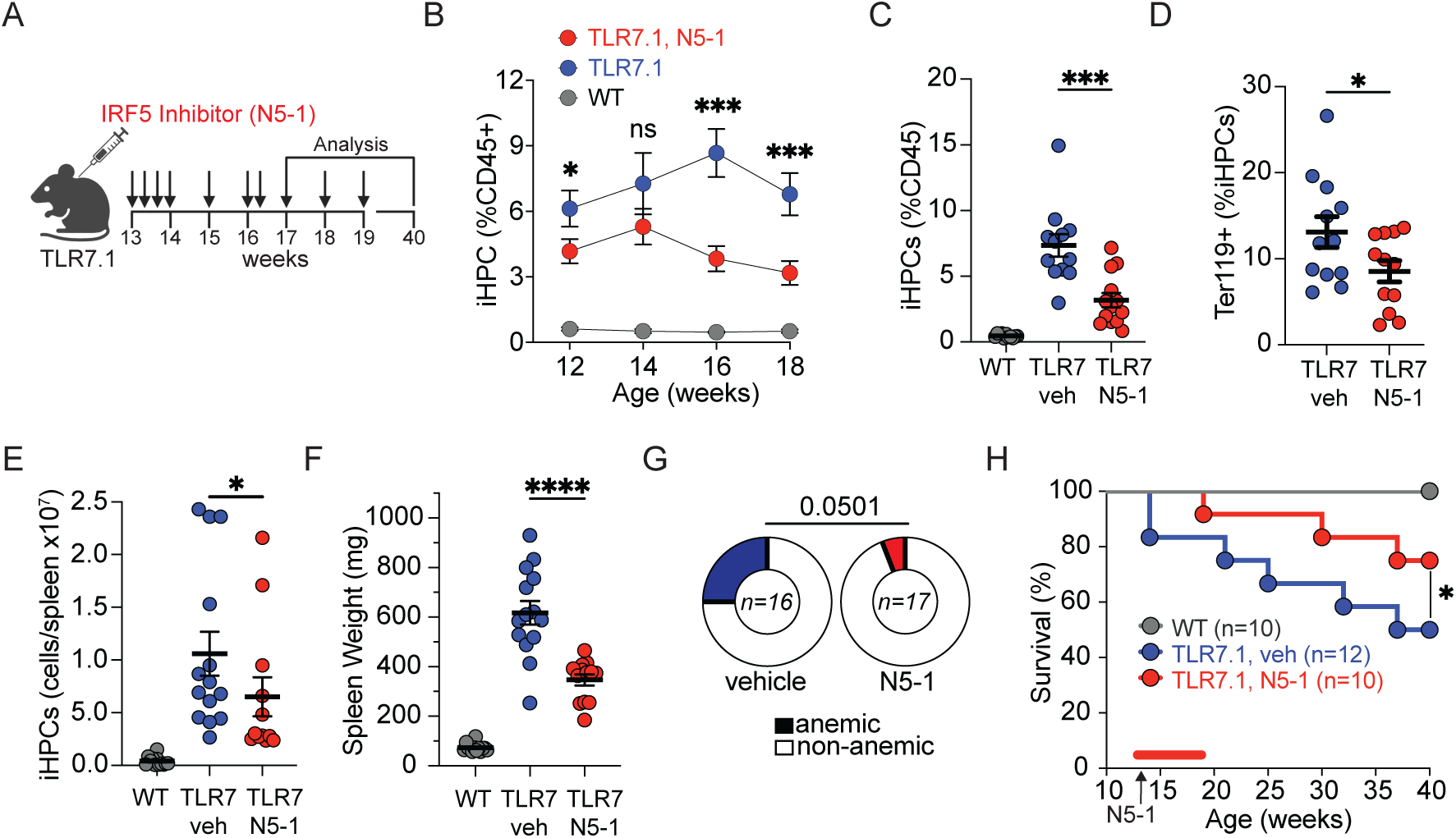
Therapeutic treatment of TLR7.1 mice with an IRF5 inhibitor reduces iHPCs and MAS-like disease. (A-E) Mice were treated with N5-1 or vehicle 10 times over 6-weeks starting at 13 weeks of age and assessed for blood iHPCs and MAS-like disease. (A) Schematic of therapeutic N5-1 treatment of TLR7.1 mice. (B-D) iHPCs and iHPC RBC phagocytosis were assessed in the blood every two weeks from 12 to 18 weeks of age (WT n=12, N5-1 treated TLR7.1 n=13, vehicle treated TLR7.1 n=13). (B) Blood iHPCs as a percentage of CD45^+^ over the course of 12 to 18 weeks. (C) Blood iHPCs as percent of CD45^+^ cells and (D) percent Ter119^+^ iHPCs at 18 weeks of age. (E-F) Mice were treated with N5-1 7 times over 4-weeks starting at 13 weeks of age and assessed for splenic iHPCs and MAS-like disease at 17 weeks of age (WT n=13, N5-1 TLR7.1 n=13, vehicle TLR7.1 n=14). (E) Splenic iHPCs and (F) spleen weights. (G) Percentage of mice that developed moderate-severe anemia (RBC <7 M/μl) in therapeutic N5-1 treated and control vehicle-treated TLR7.1 mice by 18 weeks of age (N5-1 TLR7.1 n=17, vehicle-treated TLR7.1 n=16). (H) Kaplan-Meier survival curve showing survival of WT (n=10), N5-1 treated TLR7.1 (n=12), and vehicle treated TLR7.1 mice (n=12) over 40 weeks; red bar indicates time of N5-1 treatment. *p<0.05, ***p<0.001, ****p<0.0001, Welch’s t-test (B-D); Mann-Whitney test (E-F); Wilson-Brown Binomial test (G). Log-rank (Mantel-Cox) test (H). Mean +/− SEM shown.

## DISCUSSION

We show here that cell-intrinsic IRF5 expression is required for iHPC differentiation from Ly6C^HI^ monocytes, and that myeloid IRF5 is a key contributor to MAS-like disease during TLR7-driven inflammation. Together these data position IRF5 as a regulator of both inflammation-induced monocyte differentiation and myeloid-driven inflammatory disease. In humans, SNPs associated with elevated *IRF5* expression confer genetic risk of SLE, and myeloid cells from risk-haplotype carriers show increased constitutive nuclear IRF5 translocation ^63,64^. In mouse models, B cell IRF5 is well established to drive lupus-like disease ^58–60^. Our findings extend this to IRF5 function in monocyte-derived cells, including iHPCs, and may explain the genetic risk that *IRF5* SNPs confer in MAS secondary to rheumatic disease and in secondary hemophagocytic lymphohistiocytosis due to infection ^28,29^.

Our work and prior studies support a key role for IRF5 in monocyte and macrophage differentiation during inflammation. IRF5 promotes inflammatory macrophage differentiation both in vitro, downstream of TLR4 in BM–derived macrophages, and in vivo, in models including atherosclerosis, viral infection, and *Helicobacter hepaticus*-induced colitis ^40–42,46–50^. The gene signature defining iHPCs resembles that of IRF5-regulated CD11c^+^ macrophages arising from Ly6C^HI^ monocytes in the colon during *H. hepaticus*-induced colitis^40^, sharing *Itgax* (encoding CD11c), phagocytosis genes *(C1qa-c, Fcgr4*), cytokines *(Il10, Il12b, Il1b)*, and AP-1 family members (*Maf, Batf3).* This overlap across tissues and inflammatory stimuli suggests that IRF5 drives a shared monocyte differentiation program. Despite these shared gene signatures, iHPCs are distinct from these previously described cells in that they fully depend on IRF5 for their differentiation, whereas prior studies report only partial defects in differentiation of *Irf5*-deficient inflammatory CD11c^+^ macrophages^40,41^. This difference in absolute IRF5 dependence may reflect distinct tissue environments or redundancy in signals driving these other monocyte-derived cells.

We showed that cell-intrinsic IRF5 expression is required for iHPC differentiation but not for the development of their Ly6C^HI^ monocyte progenitors. Both germline and myeloid-specific *Irf5* deletion reversed the monocytosis in TLR7.1 mice, which we previously showed results from emergency myelopoiesis^25,26^. Therefore, IRF5 promotes emergency myelopoiesis in a monocyte extrinsic manner, consistent with our earlier finding that TLR7-driven iHPC differentiation requires cell-intrinsic *Tlr7* expression while Ly6C^HI^ monocyte expansion does not^15^. Reduced monocytosis in *Irf5*^−/−^ mice during chikungunya virus infection and *H. hepaticus*-induced colitis is consistent with a broader model whereby cell-extrinsic IRF5 drives emergency myelopoiesis across inflammatory contexts^40,46^. This indirect effect of IRF5 in monocytosis is likely due to a role in inducing cytokines that act on hematopoietic stem and progenitor cells, and it is intriguing to speculate that iHPCs may contribute cytokines for this emergency myelopoiesis. This indirect role in myelopoiesis is distinct from the cell-intrinsic requirement for IRF5 in iHPC differentiation and CD11c^+^ macrophage development during *H. hepaticus*-induced colitis^40^.

IRF5 is critical for lupus-like disease development in several mouse models and in some cases monoallelic *Irf5* deletion is sufficient to ameliorate disease^35,58,60,65–67^, similar to what we show here in TLR7.1 mice, where monoallelic *Irf5* deletion fully protects not only against lupus-like disease, but also MAS. Previous work in lupus models has focused on the importance of *Irf5* in B cells ^58,67^, with *Irf5* deletion in B cells, but not in myeloid cells, sufficient to confer full protection in the *FcγRIIB*^−/−^*Yaa* lupus model^59^. In contrast, in TLR7.1 mice, myeloid-specific loss of *Irf5* protected against both MAS-like and lupus-like disease. This is the first demonstration that myeloid-intrinsic *Irf5* deficiency alone is sufficient to protect against either disease. In TLR7.1 *Irf5*^ΔM^ mice, the dramatic reduction in iHPCs suggests that loss of this population contributes to protection. TLR7.1 *Irf5*^ΔM^ mice also had reduced serum anti-RNA autoantibodies, indicating reduced pathogenic B cell activity, even though *Irf5* expression is intact in B cells, and plasma cell and GC B cell expansion persist in these mice. We therefore propose that reduced iHPC accumulation dampens systemic inflammation, indirectly limiting B cell activation. Additionally, although cell-intrinsic IRF5 expression was not required for neutrophil development in TLR7.1 mice, loss of IRF5 in neutrophils may also contribute to disease protection. Together, these findings identify a myeloid-intrinsic, B cell-independent, role for IRF5 in driving MAS-like and lupus-like disease.

Our data support a model in which IRF5 establishes iHPC identity through regulation of a functional iHPC gene program, possibly acting with other transcription factors at accessible chromatin. Relative to all upregulated iHPC genes, those associated with IRF5-binding accessible chromatin showed higher enrichment for key iHPC-specific gene programs, consistent with IRF5-binding genes constituting the core iHPC transcriptional identity. Several IRF5-dependent genes induced in vitro during the Ly6C^HI^ monocyte-to-HPC transition, and that are upregulated in iHPCs in vivo, including *Tnfaip3, C1qa/C1qb, Ifih1, Il10, Isg15,* and *Nfkbia,* are orthologs of human genes implicated in autoimmune or autoinflammatory disease ^68^. This mix of proinflammatory genes and negative regulators of NF-κB signaling suggests that IRF5 regulates both positive and negative arms of the inflammatory response in iHPCs. Indeed, NF-κB signaling was among the most enriched pathways upregulated in iHPCs, and this enrichment increased when analysis was limited to known IRF5-binding regions. Additional enriched pathways included cytokine production/signaling and phagocytosis-related programs, including complement opsonization (*C1qa-c*), iron handling (*Cp*), and FγcR-mediated phagocytosis. Together, these implicate IRF5 in the regulation of phagocytosis and iron-handling machinery in iHPCs. In contrast, IRF5-binding accessible regions in TLR7.1 Ly6C^HI^ monocyte upregulated genes were enriched for few pathways beyond interferon signaling, and ISGs were unchanged between TLR7.1 and TLR7.1 *Irf5*^−/−^ Ly6C^HI^ monocytes. Redundancy among IRF5, IRF3, and IRF7 in myeloid type I IFN and ISG responses may allow this program to remain intact without IRF5 ^69^. Together with the low nuclear IRF5 signal in these cells, this argues against direct regulation of the monocyte transcriptional program by IRF5. This contrasts with the non-redundant requirement for IRF5 in the iHPC program, highlighted by the loss of iHPCs in its absence.

IRFs orchestrate inflammatory responses and immune cell development by binding to specific DNA motifs as homo- or heterodimers. IRF dimer composition and transcription factor cofactor interactions determine DNA binding affinity and motif specificity, thereby shaping gene expression and chromatin remodeling. IRF composite motifs are well established in myeloid cell identity; for example, IRF8-AP1 and IRF8-PU.1 heterodimers cooperate to establish and maintain the cDC1 gene expression program^43,44,70,71^. Overlapping our accessible chromatin peaks with known IRF5 binding sites from TLR9-activated cDC1s^51^ revealed distinct IRF motif enrichment in IRF5-binding regions of iHPCs versus TLR7.1 Ly6C^HI^ monocytes. This finding, together with the increased expression in iHPCs of specific ETS and AP-1 family members such as *Batf* and *Ets2*, suggests that IRFs drive cell-specific gene programs through distinct combinations of IRF dimers and cofactors. During in vitro cultures modeling early iHPC differentiation, we found that TLR7 signaling upregulated *Batf* and *Ets2* in Ly6C^HI^ monocytes in an IRF5-dependent manner, supporting a functional link between IRF5 and these transcription factors during iHPC differentiation. Our data also suggest that IRF5 and NF-κB subunits may cooperate in cis to regulate iHPC gene programs downstream of TLR7 signaling, with an enrichment of NF-κB motifs in iHPC-specific IRF5-binding regions of open chromatin, consistent with previous work showing that IRF5 can regulate inflammatory gene expression in concert with the NF-κB subunit RelA in TLR4-stimulated BMDMs^39^. This cooperation between IRF5 and NF-κB may be particularly important downstream of TLR7, where both transcription factors are activated in parallel. Together, our data support a model in which IRF5 drives iHPC-specific gene programs through combinatorial interactions with other IRFs, NF-κB, ETS, and AP-1 transcription factors.

As mortality in patients who develop MAS ranges from 5-30%^6,72^, effective therapies are essential. Current therapies dampen overall immune responses with high-dose systemic steroids and immunosuppressants, as well as more targeted biologics, such as the IL-1 receptor antagonist anakinra, the anti-IFNγ monoclonal antibody emapalumab, and JAK inhibitors^53,72^. However, these therapies are not effective for all patients, and MAS can still occur despite adequate control of the underlying rheumatic disease. IRF5 inhibition is therefore a compelling therapeutic target for both MAS and for certain underlying rheumatic diseases, including SLE. We and others have shown in multiple murine lupus models that IRF5 inhibitor treatment as well as *Irf5* deletion, even after disease onset^73^, reduces lupus-like disease and disease-associated immune cell activation^62,74^. Here we showed that therapeutic IRF5 inhibition reduces MAS features and underlying lupus-like disease in TLR7.1 mice, suggesting this may be a potential therapeutic strategy for MAS.

Our study has several limitations. Although we show a clear role for IRF5 in iHPC differentiation, without a model for iHPC-specific *Irf5* deletion we cannot disentangle the impact of IRF5 on iHPC differentiation from its impact on function, an important consideration given that other IRFs regulate both processes. IRF8, for example, governs cDC1 differentiation and identity while also regulating effector functions, such as cross-presentation^43,44,70,71^. Additionally, while we were able to identify accessible peaks with confirmed IRF5 binding in endosomal TLR-stimulated myeloid cells by intersecting our ATAC-Seq data with a publicly available IRF5 ChIP-seq dataset from TLR9-stimulated dendritic cells^51^, our interpretations are limited because the chromatin landscapes of dendritic cells, monocytes, and iHPCs are not identical. Therefore, our analysis may have missed important IRF5-binding regions specific to iHPCs, including in genes underlying iHPC-specific functions such as RBC phagocytosis and degradation. Mapping IRF5-binding regions within the native iHPC chromatin landscape is an important next step.

Although extensive literature implicates IRF5 in inflammatory macrophage differentiation, the in vivo mechanisms governing this process remain unclear. Our work demonstrates a critical, cell-intrinsic requirement for IRF5 in a specific Ly6C^HI^ monocyte-derived macrophage subset that drives pathology during chronic TLR7 signaling. Together with the efficacy of therapeutic IRF5 inhibition in TLR7.1 mice, these findings identify IRF5 as a compelling target in MAS and begin to define how IRF5 acts with other transcription factors to regulate inflammatory macrophage development.

## METHODS

### Mice

TLR7.1 mice were obtained from Dr. Sylvia Bolland, NIH (MGI:5444399)^14^. *Irf5^−/−^* mice were originally obtained from Dr. Tadatsugu Taniguchi via the Rifkin Lab and backcrossed 11 generations to the C57BL/6 background (MGI:3576384)^33^. B6.129P2-*Lyz2*^tm1(cre)Ifo^/J (*Lyz2*-Cre; JAX stock #004781), C57BL/6J (JAX stock #000664), B6.SJL-*Ptprc*^a^ *Pepc*^b^/BoyJ (B6.SJ; JAX stock #002014), C57BL/6J-*Ptprc*^em6Lutzy^/J (B6.CD45.1; JAX stock #033076), and C57BL/6-*Irf5*^tm1Ppr^/J (*Irf5*^FL/FL^; JAX stock #017311) mice were obtained from Jackson Labs. TLR7.1, TLR7.1 *Irf5*^−/−^, TLR7.1 *Irf5*^+/−^, TLR7.1 *Lyz2*^CRE/CRE^, and TLR7.1 *Lyz2*^CRE/CRE^ *Irf5*^FL/FL^ mice were maintained on a C57BL/6 background. All experiments were performed under approved protocols from the Benaroya Research Institute or Feinstein Institute Institutional Animal Care and Use Committee.

### BM Chimeras

Mixed BM chimeras were generated by lethally irradiating (1,000 rad) recipient mice and reconstituting them with a total of 1×10^7^ BM cells. For TLR7.1:TLR7.1 and TLR7.1:TLR7.1 *Irf5*^−/−^ mixed BM chimeras in Fig. 6, B6.SJL recipient mice were reconstituting with a 1:1 ratio of TLR7.1×B6.SJL F1 and either TLR7.1 or TLR7.1 *Irf5*^−/−^ BM cells. For paired ATAC seq and RNA seq TLR7.1:TLR7.1 and TLR7.1:TLR7.1 *Irf5*^−/−^ mixed BM chimeras, B6.CD45.1 recipient mice were reconstituting with a 1:1 ratio of TLR7.1xB6.CD45.1 F1 and either TLR7.1 or TLR7.1 *Irf5*^−/−^ BM cells. For WT:WT and WT: *Irf5*^−/−^ mixed BM chimeras, recipient C57BL/6xB6.CD45.1 F1 mice were reconstituted with a 1:1 ratio of B6.CD45.1 and either C57BL/6 or *Irf5*^−/−^ BM cells. Mice were euthanized 7-20 weeks post reconstitution.

### In vivo IRF5 inhibitor treatment

IRF5 peptide inhibitor N5-1 was synthesized by LifeTein, LLC and purity was confirmed by HPLC and mass spectrometry^62^. N5-1 was evaluated in both preclinical and therapeutic treatment regimens. For both approaches, male and female WT and TLR7.1 mice received a total of 10 intraperitoneal (i.p.) injections over 4 weeks. Vehicle control groups were administered PBS, while experimental groups received 100 µg/mouse of N5-1 per injection. The injection schedule spanned 42 days (on days 0, 1, 4, 7, 14, 15, 21, 28, 35, and 42). Preclinical treatment began 5 weeks of age; therapeutic treatment started at 13 weeks of age. Mice in the therapeutic treatment cohort evaluated at 17 weeks of age received 7 N5-1 treatments over 4 weeks.

### Hematology monitoring and kidney pathology

To assess anemia and thrombocytopenia, mice were bled retroorbitally with heparinized tubes or, for the IRF5 inhibitor experiments, submandibularly and collected into EDTA tubes. Blood was analyzed on the Hemavet® Hematology Analyzer (Drew Scientific). Kidneys were fixed in neutral buffered formalin and paraffin embedded. Paraffin-embedded kidney sections were stained with Periodic Acid–Schiff (PAS). Histopathologic features of glomerular lesions were graded for severity in a double-blinded manner. Disease features scored included: Mesangial hypercellularity, Mesangial “hyaline” expansion, Endocapillary hypercellularity, Hyalinization (capillary wall hyaline deposits), Crescents (cellular and fibrocellular), and Glomerulosclerosis. The glomerular lesions per kidney were scored as follows: 0, absent; 1+ (1-10% of glomeruli affected); 2+ (11-25% of glomeruli affected); 3+ (26-49% of glomeruli affected); and 4+ (greater than 50% of glomeruli affected). The total Disease Score was calculated as the sum of all disease feature scores.

### Cell isolation, Flow Cytometry, and Cell Sorting

Single cell suspensions were prepared from mouse peripheral blood or spleen. For iHPCs and monocytes, spleens were digested in a cocktail of 0.17 mg/mL Liberase TL (Sigma, #5401020001) and 40 μg/mL DNase1 (Sigma, #11284932001) in complete RPMI (Hyclone) as previously described^15^. For B and T cells, spleens were mechanically disrupted in complete RPMI. Erythrocytes were removed from peripheral blood and splenocytes by ACK Lysis Buffer (Biolegend, #420302; Lonza, #Bp10-548e). Non-specific binding of antibodies to Fc-receptors was blocked with 65 μg/mL each polyclonal rat and mouse IgG (Sigma, #I8015 and #I8765) or with anti-CD16/32 antibody before cell surface staining with fluorescently labeled antibodies. All antibodies used for flow cytometry can be found in Tabel S1. Cells were then washed and, for some panels, stained with fixable viability dye (Fisher scientific) in PBS. Samples were washed and fixed with Fixation and Permeabilization buffer (BD Biosciences, #BDB554714). For samples with quantification of RBC phagocytosis, 71.4 μg/ml purified anti-mouse TER-119 (Biolegend, #116202) was included in the pre-fix block. Post fix, samples were washed in BD Perm/Wash buffer, blocked in 65 μg/mL polyclonal rat IgG, and stained with fluorescently labeled anti-Ter-119 to detect intracellular RBCs. To accurately assess phagocytosis, some samples were blocked intracellularly with 25 μg/ml purified anti-mouse Ter-119 prior to intracellular staining with fluorescently labeled anti-Ter-119. Data were acquired on an LSRII, FACS Symphony, or LSR Fortessa (BD Biosciences), and analyzed using FlowJo (BD Biosciences). Cell yield was quantified using polystyrene microspheres (Polysciences).

Cell sorting was performed on a FACSAria (BD Biosciences). For real-time PCR and ATAC sequencing, samples were sorted into complete RPMI, for Bulk RNA sequencing cells were sorted directly into reaction buffer from the SMART-Seq v4 Ultra Low Input RNA Kit for Sequencing (Takara). To verify the purity of the sorted populations, all the cell sorting steps were validated in post-sort analysis.

### Imaging flow cytometry

Imaging flow cytometry was performed on the Amnis Imagestream. Briefly, mouse PBMC or splenocytes were isolated followed by extracellular staining indicated in Table S1. After surface staining, cells were fixed in 4% PFA for one hour at room temperature, followed by permeabilization overnight with 0.5% Triton X-100 in 5% BSA. Permeabilized cells were blocked in 5% BSA solution and stained for intracellular IRF5 (Abcam, clone #EPR17067) and an anti-Rabbit APC Secondary (Thermo Fisher Scientific), DAPI was used for nuclear staining (Thermo Fisher Scientific).

### In Vitro Cell Culture

BM Ly6C^HI^ monocytes were isolated by negative selection using the mouse Monocyte Isolation Kit (Miltenyi, #130-100-629) per manufacturer’s protocol. 40,000 cells, for flow cytometry, or 100,000 cells, for RNA sequencing, were plated per well in serum-free RPMI for 30 minutes after which media was replaced with complete RPMI and the cells were rested for 2 hours. Cells were cultured in 10 ng/ml of M-CSF (Peprotech, #315-02) with or without 1 μg/ml of R848 (Invivogen, tlr7-r848-5). For flow cytometric quantification of CD11b^+^CD31^+^ cells, 48 hours after the addition of R848, adherent cells were washed with 1x PBS and dissociated from the plate with cell dissociation buffer (Life Technologies). As described above, cells were blocked with rat and mouse IgG, stained with as indicated in Table S1, and analyzed by flow cytometry.

### Quantitative RT-PCR

Sorted cells were lysed in RLT plus lysis buffer (Qiagen) with β-mercaptoethanol. RNA was generated using RNeasy Plus Micro Kit (Qiagen). cDNA was synthesized using Primescript (Takara), random hexamers (Thermo Scientific), and OligoDT primers (Thermo Scientific). qPCR was performed using SYBR green reagents (Takara) on a QuantStudio 5 Real-Time PCR System (Thermofisher). Arbitrary units were calculated using the ΔΔCT method normalized to *Hprt*.

### Autoantibody Analysis

Anti-RNA IgG was measured by ELISA. Plates were pretreated with 0.005% poly-L-lysine, coated with 100 µg/mL baker’s yeast RNA (Sigma, #R6750) overnight at 37°C, and blocked with 2% BSA/PBS. Plasma (1:100) was incubated overnight at 4°C, followed by alkaline phosphatase-conjugated goat anti-mouse IgG (Southern Biotech, #1030-04). ELISA was developed with PNPP and absorbance read at 450 nm.

### Bulk RNA-Seq and Analysis

For R848-treated WT vs *Irf5*^−/−^ Ly6C^HI^ monocytes, 100,000 BM Ly6C^HI^ monocytes were cultured, as described above. Cells were lysed in RLT plus lysis buffer (Qiagen) and RNA generated using the RNeasy micro kit (Qiagen), followed by reverse transcription and PCR amplification to generate full-length amplified cDNA. For iHPCs and monocytes from BM chimeras, 600 cells were sorted directly into reaction buffer from the SMART-Seq v4 Ultra Low Input RNA Kit for Sequencing (Takara) and snap-frozen. RNA libraries were built using SMART-Seq v4 Ultra Low Input RNA Kit for Sequencing (Takara Bio USA, Cat #634891) following manufacturer’s instructions. Sequencing libraries were constructed using the Nextera XT DNA Library Preparation Kit with unique dual indexes (Illumina) to generate Illumina-compatible barcoded libraries. Libraries were pooled and quantified using a Qubit® Fluorometer (Life Technologies). Sequencing of pooled libraries was carried out on a NextSeq 2000 sequencer (Illumina) with paired-end 59-base reads, using NextSeq P2 sequencing kits (Illumina) with a target depth of 5 million reads per sample. Base calls were processed to FASTQs on BaseSpace (Illumina), and a base call quality-trimming step was applied to remove low-confidence base calls from the ends of reads using fastp (0.23.1). Trimmed reads were aligned using STAR aligner (v2.7.11b)^75^ with the Genome Reference Consortium mouse genome assembly (GRCm38) and gene annotations from Ensembl release 91^76^. Gene counts were generated using HTSeq-count (v 2.0.2)^77^. Quality control metrics were calculated using Picard (v 3.1.0)^78^, FastQC (v0.12.1)^79^, and Samtools (v1.16)^80^. Samples that passed quality criteria (FASTQ reads > 1 million, mapped reads > 70%, and median CV coverage < 0.9) were retained for analysis.

Bulk RNA-Seq analysis was performed in R (v4.5.2). Protein-coding genes with expression of at least 1 CPM in 10% of libraries were used in the analysis. Expression counts were filtered and normalized using the TMM algorithm^81^, and principal component analysis (PCA) was performed on log2-transformed TMM-normalized counts. Differential expression analysis was performed using the linear models for microarray data (limma) package^82^. The limma voomWithQualityWeights transformation of TMM-normalized counts was used to fit a linear model of gene expression with mouse identity as a random effect. A false discovery rate (FDR) adjustment was applied to correct for multiple testing. Differentially expressed genes (DEGs) were defined as those with an absolute log2 fold change ≥ 1 and Benjamini-Hochberg-adjusted p-value < 0.05. For the RNA-seq of BM Ly6C^HI^ monocytes stimulated with R848, the linear model of gene expression was a function of genotype and time post R848-treatment. To compare the transcriptional response to R848 stimulation between genotypes, scatter plots were generated where the log2 fold change of R848-treated versus untreated cells at each timepoint was plotted for WT cells (x-axis) against IRF5 KO cells (y-axis). Genes were classified as upregulated in WT cells only, IRF5 KO cells only, both genotypes, or neither (absolute log2 fold change ≥ 1, FDR < 0.05).

### Omni-ATAC Seq and Analysis

Splenic cells were isolated and stained as described above. iHPCs, Ly6C^HI^ monocytes, and patrolling monocytes were sorted into cold complete RPMI, cells were counted, and 20,000-50,000 cells were resuspended in cold 1x PBS. Nuclei were isolated and tagmented using a previously published Omni-ATAC protocol^83,84^. Transposed DNA was purified using the QIAGEN MinElute Reaction Cleanup kit (28204) and eluted in 12 μl of elution buffer (10 mM Tris-HCl, pH 8). Transposed fragments were amplified with custom Nextera primers for amplifying libraries (Nextera_F AATGATACGGCGACCACCGAGATCTACACXXXXXXXXTCGTCGGCAGCGTC; Nextera_R CAAGCAGAAGACGGCATACGAGATXXXXXXXXGTCTCGTGGGCTCGG) with indices and sequencing adapters as described^85^. Libraries were purified using a 0.5x/1.5x double-sided selection with AMPure XP beads (Beckman Coulter) per manufacturer’s protocol. Libraries were multiplexed at an equimolar concentration as determined by sequencing a small portion of each library on a MiSeq (Ilumina MiSeq Reagent Kits V2 500 cycles MS-102-2003). Pooled libraries were quantified by Qubit and Tape Station before dilution and sequencing. All Omni-ATAC libraries were sequenced using paired-end, dual-index sequencing on a NextSeq instrument.

Adapter sequences were trimmed from paired-end ATAC-seq reads using Trimmomatic (v0.39; NexteraPE-PE.fa:2:30:10 LEADING:3 SLIDINGWINDOW:4:20 MINLEN:8)^86^, then mapped to the genome (GRCm39) using STAR (v2.7.11b)^75^. Reads mapping to mitochondrial DNA were removed using SAMtools (v1.20)^80^, and duplicate reads were removed using Picard (v3.3.0; MarkDuplicates REMOVE_DUPLICATES=true)^78^. Peaks were called using MACS (v2.2.9.1; -f BAMPE -q 0.001)^87^. Sample quality was assessed using enrichment of reads around transcription start sites (TSSe) and fractions of reads in peaks (FRiP), and samples passing quality criteria (FASTQ reads > 30 million, TSSe scores > 10, FRiP score > 0.2) were retained for analysis.

Differential chromatin accessibility was analyzed with the DiffBind package (v3.16.0)^88^ in R. Peaks smaller than 50 bp, greater than 10000 bp, or falling within 1Mb of *Irf5* or the TLR7.1 BAC regions were excluded from analysis, as were peaks falling within the GRCm39 exclusion regions included in the excluderanges package (v0.99.8)^89^. For each pairwise differential chromatin accessibility comparison (iHPCs versus TLR7.1 Ly6C^HI^ monocytes, TLR7.1 *Irf5*^−/−^ Ly6C^HI^ monocytes versus TLR7.1 Ly6C^HI^ monocytes), consensus peaks were counted and recentered using the dba.count() function with peak summits recentered including 250 bp upstream and downstream of the original peak summit. Samples were normalized using the DiffBind settings library = DBA_LIBSIZE_FULL, normalize = DBA_NORM_LIB, background = FALSE, and differential accessibility was analyzed with DiffBind using EdgeR. Significant differentially accessible regions (DARs) were defined as those with absolute log2 fold change >= 1 and FDR-adjusted p-value >= 0.05 and annotated to the nearest gene bodies using ChIPpeakAnno (v3.40.0)^90^. DARs were overlapped with IRF5 ChIP peaks using BEDTools intersect (v2.31.1)^91^.

HOMER findMotifsGenome.pl (v5.1; -size 200 -mask)^92^ was used to find enrichment of known motifs in DARs. For DARs without IRF5 ChIP overlap, random genome regions with matched GC content were used as background for motif finding, while for DARs with IRF5 ChIP overlap, non-IRF5-overlapping DARs with matched GC content were used as background.

Motif similarities were calculated using the compare_motifs() function of the universalmotif package in R (v1.24.2; method=“PCC”, min.overlap=5)^93^.

### AI usage statement

Claude Code (Anthropic, Sonnet 5) was used to generate code and plots for some figures; all AI-generated code was reviewed by a bioinformatician at the Benaroya Research Institute. Specifically, Claude Code was used to generate and run over-representation analysis (ORA), to generate lollipop plots of non-redundant enriched pathways (Fig. 1 D-E, H-I), to generate the motif similarity heatmap (Fig. S2 B), the z-scored motif heatmap (Fig. S3 D), and gene tracks for select genes (Fig. 2 D, Fig. S3 E).

In the initial editing process Microsoft 365 Copilot (Microsoft, 2026) and Claude (Anthropic, Sonnet 5) were used in select sections of the manuscript text to provide suggestions for sentence structure and clarity. All suggested changes were reviewed carefully for scientific accuracy and, in some instances, were integrated into the text.

### Statistical analysis

Normality of data distribution was assessed using the Shapiro-Wilk test and equality of variances was assessed using the F-test for two groups and the Brown-Forsythe test for 3 or more groups. For comparisons of two groups, a Student’s unpaired t-test was used when variances were equal, and a Welch’s t-test was used when variances were unequal. For comparisons of three or more groups with equal variances, ordinary one-way ANOVA was used followed by a Dunnett’s multiple comparisons test (for comparisons to a single control) or Šídák’s multiple comparisons test (for pairwise comparisons between selected groups). When variances were unequal, Brown-Forsythe and Welch ANOVA was used followed by the Dunnett’s T3 multiple comparisons test. For data sets that did not follow a normal distribution, the Mann-Whitney test was used for comparisons of two groups, and the Kruskal-Wallis test followed by Dunn’s multiple comparisons test was used for comparisons of three or more groups. All tests were two-tailed, with a significance set at α = 0.05. Time spent anemia-free and thrombocytopenia-free as well as survival data were analyzed using the Log-rank (Mantel-cox) test, binomial proportions were compared using the Wilson/Brown method for confidence intervals. Statistical analysis for parametric, non-parametric, and survival comparisons was performed in GraphPad Prism (Prism Version 11, GraphPad Software, Boston, MA). Statistical significance of the overlap between IRF5-dependent R848-induced genes and upregulated iHPC genes was assessed using a one-tailed exact hypergeometric test, with background set as the total number of genes (12,473) present in both data sets (phyper function, R v[4.4.1]). Tests used for each specific comparison are indicated in the corresponding figure legends or methods.

### Data and code availability

Analyzed bulk RNA seq and ATAC seq data are presented in the Extended Data files. FASTQ files for the bulk RNA-Seq and BED files for the ATAC-Seq will be uploaded to GEO. Code will be available at https://github.com/BenaroyaResearch. Data and code deposition are in progress.

## Acknowledgements

We thank Jenna Battaglia, Lucy Li and Dr. Minjian Ni for help with experiments, Dr. Susan Canny for insights about MAS, Dr. Alanna Sholokhova and Dr. Andrew Koval for review and discussion of statistical tests, Dr. Hannah DeBerg for reviewing code and for advice on analysis strategies, Stephanie Ryder for sharing Omni-ATAC methodology, and Kimberly O’Brien, Vivian Gersuk and Basilin Benson in the Benaroya Research Institute Genomics Core. We acknowledge the BRI Animal Resources Core, the BRI Cell and Tissue Analysis Core (RRID: SCR_026327) for flow cytometry and histology, the BRI Genomics Core (RRID: SCR_026658) for RNA-seq library generation and sequencing and ATAC-seq sequencing. We also thank the M.J. Murdock Charitable Trust for generously providing equipment funding for the BRI Cores. All schematics were created using Biorender.com.

---

**Fig. S1:**
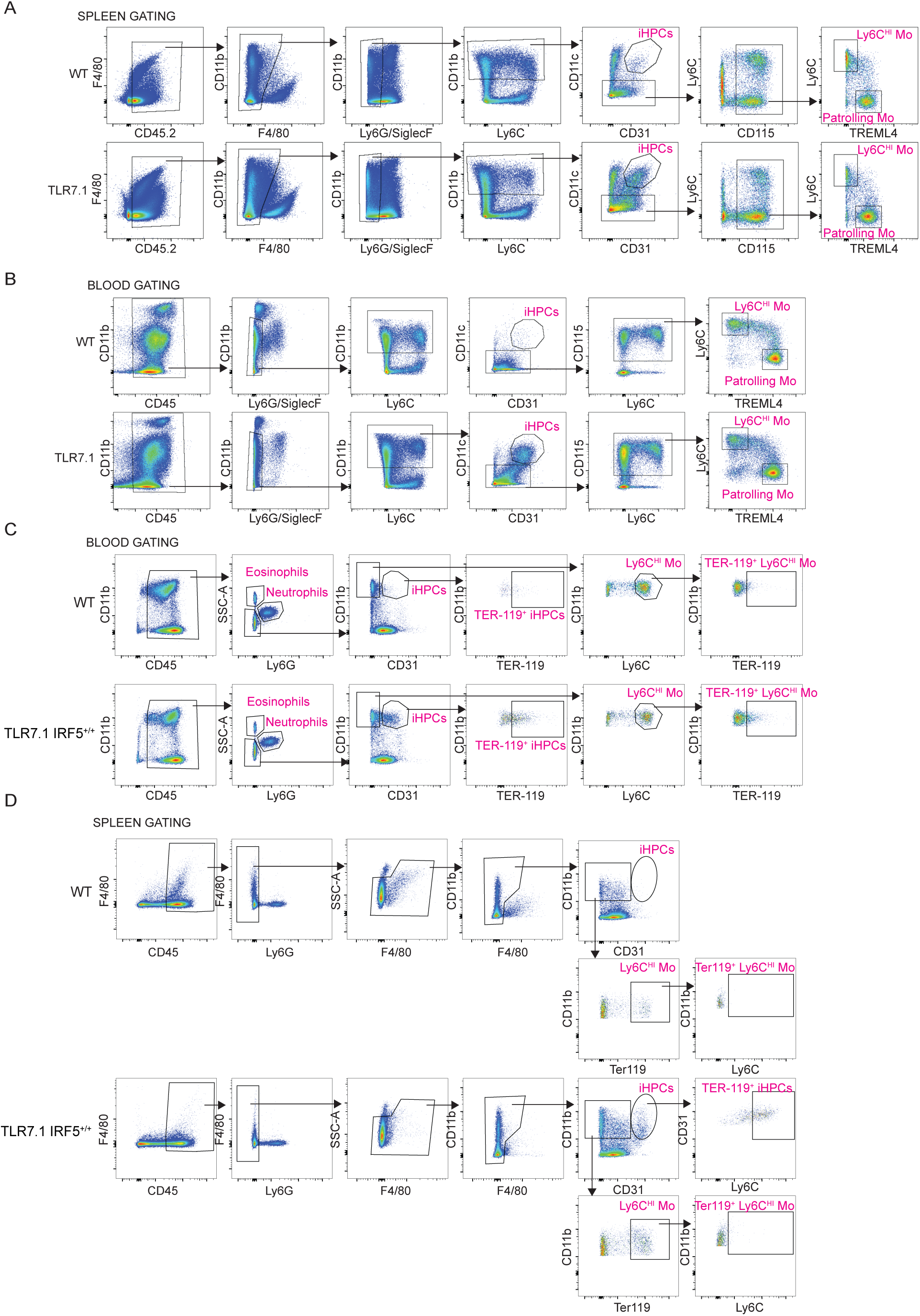
iHPC and Ly6C^HI^ monocyte gating. (C) Gating scheme for iHPCs, Ly6C^HI^ monocytes, and patrolling monocytes staining in the spleen (A) and blood (B) from TLR7 and WT chimeras used in Fig. 1 B, Fig. 4, Fig. 5 (no CD115 staining), Fig. S5, Fig. S6. Full gating scheme for iHPCs, Ly6C^HI^ monocytes, and TER-119^+^ intracellular staining in the blood (C) and spleen (D) of TLR7.1 and WT mice used in Fig. 1 K-L, Fig. 3, Fig. 6, Fig. S4, Fig. S7.

**Fig. S2:**
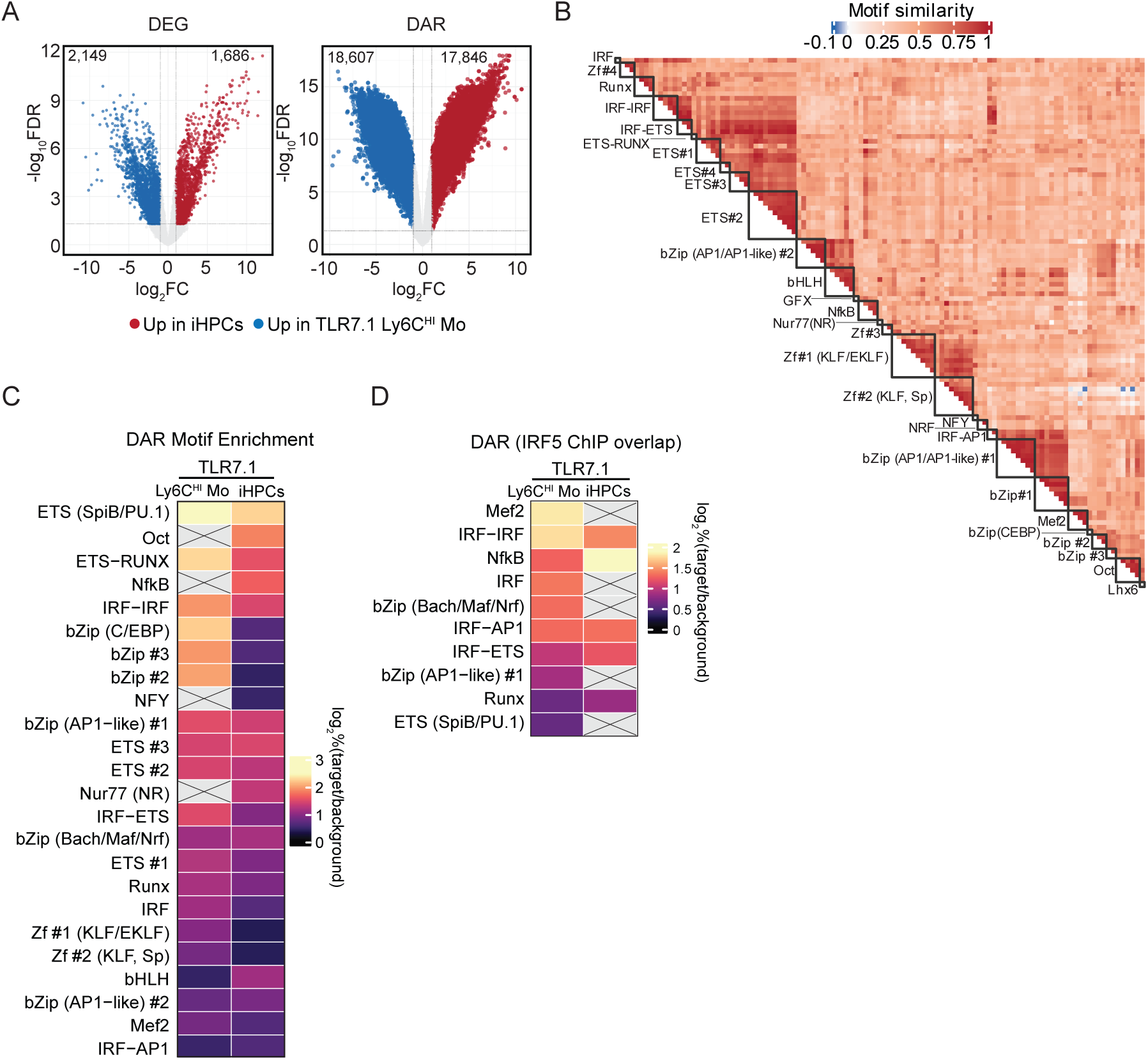
IRF5 drives an iHPC transcriptional program distinct from Ly6C^HI^ monocytes. (A-D) Paired bulk RNA sequencing and ATAC sequencing of iHPCs (n=4), TLR7.1 Ly6C^HI^ monocytes (n=4-6) sorted from TLR7.1:TLR7.1 chimeras and WT:WT chimeras (A) Volcano plots showing the differential gene expression (DEG, left) and chromatin differential accessible regions (DAR, right) between TLR7.1 Ly6C^HI^ and iHPCs (log_2_Fc ≥ 1, adj.p <0.05). Increased expression or accessibility in TLR7.1 Ly6C^HI^ monocytes in red; increased expression or accessibility in iHPCs in blue. (B) Similarity heatmap of significantly enriched motifs (p < 1e-40) in TLR7.1 Ly6C^HI^ monocytes and iHPCs from HOMER known motif analysis. Similar motifs are labeled as indicated on left.(C) Heatmap of all significantly enriched HJOMER known motif families in the differentially accessible peaks between TLR7.1 Ly6C^HI^ monocytes and iHPCs, shown as frequency of the motif within open peaks (target) relative to frequency of the motif within all accessible peaks normalized for GC content (background). (D) Heatmap of all HOMER known motif enrichment of all enriched motif families with published IRF5 binding in the differentially accessibly peaks between TLR7.1 Ly6C^HI^ monocytes and iHPCs, indicated as frequency of the motif within open peaks with known IRF5 binding (target) relative to frequency of the motif within non-IRF5-bound accessible peaks normalized for GC content (background).

**Fig. S3.**
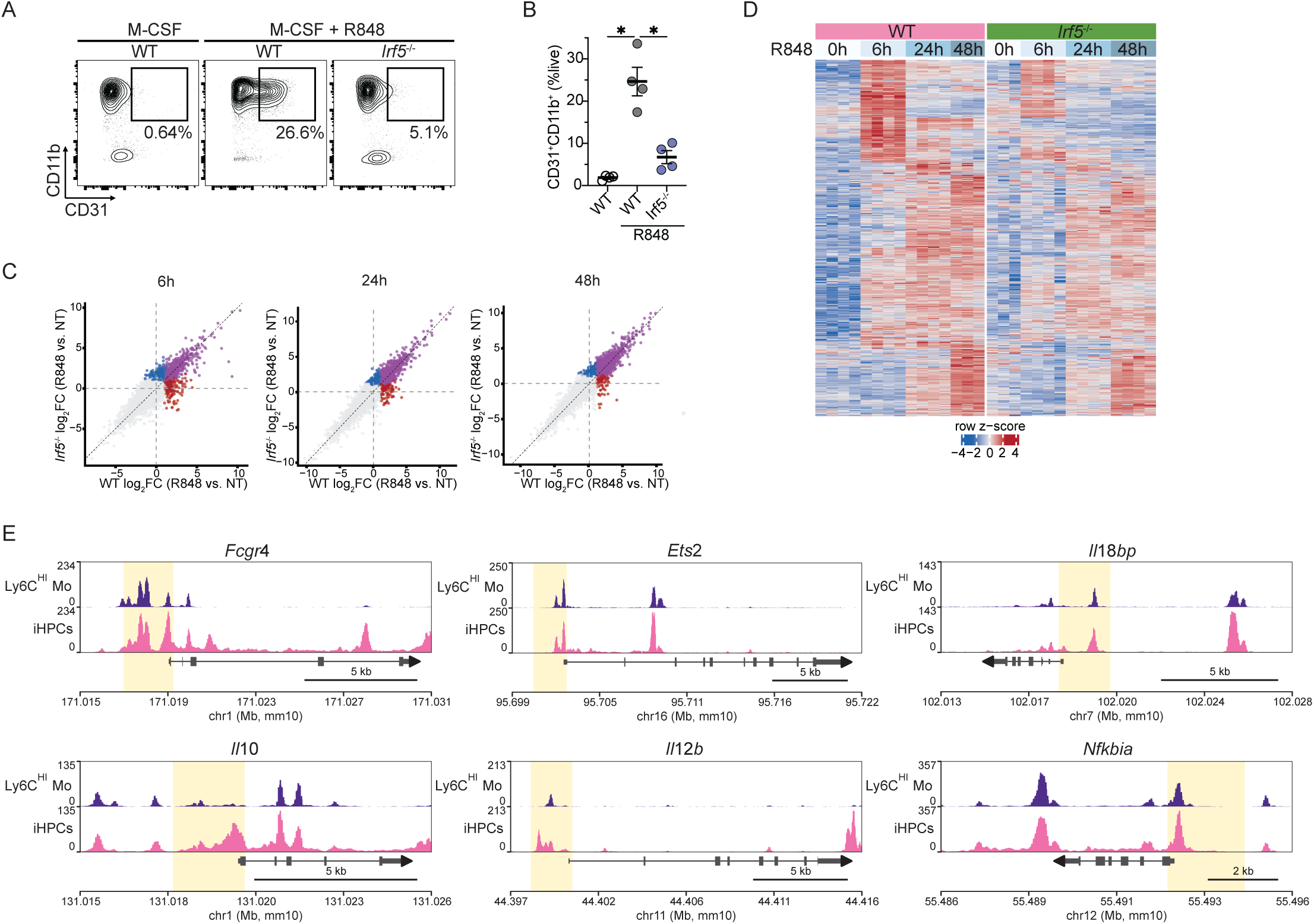
R848 induces an IRF5-dependent gene program in Ly6C^HI^ monocytes. (A, B) WT and *Irf5^−/−^*BM Ly6C^HI^ monocytes were treated for 48 hours with M-CSF +/− R848. Representative flow cytometry (A) and quantification (B) of CD11b^+^CD31^+^ iHPC-like cells as a percent of live cells (n=4 per group). (C) Bulk RNA sequencing of WT and *Irf5*^−/−^ Ly6C^HI^ monocytes at baseline (0h, NT) and with 6, 24, and 48 hours of R848 and M-CSF. XY correlation plots between R848 induced genes in WT and *Irf5*^−/−^ Ly6C^HI^ monocytes at 6, 24, or 48 hours. Genes significantly induced by R848 2-fold in both *Irf5*^−/−^ and WT Ly6C^HI^ monocytes are shown in purple, genes selectively significantly upregulated in *Irf5*^−/−^ or WT Ly6C^HI^ monocytes are shown in blue and red, respectively. (D) Heatmap of IRF5-dependent genes that were significantly induced 2-fold over the 0 hour timepoint by R848 in WT Ly6C^HI^ monocytes and significantly less induced in *Irf5^−/−^* cells, as shown in Fig. S2 C. (E) ATAC-seq tracks for select genes from Fig. 2 C.

**Fig. S4.**
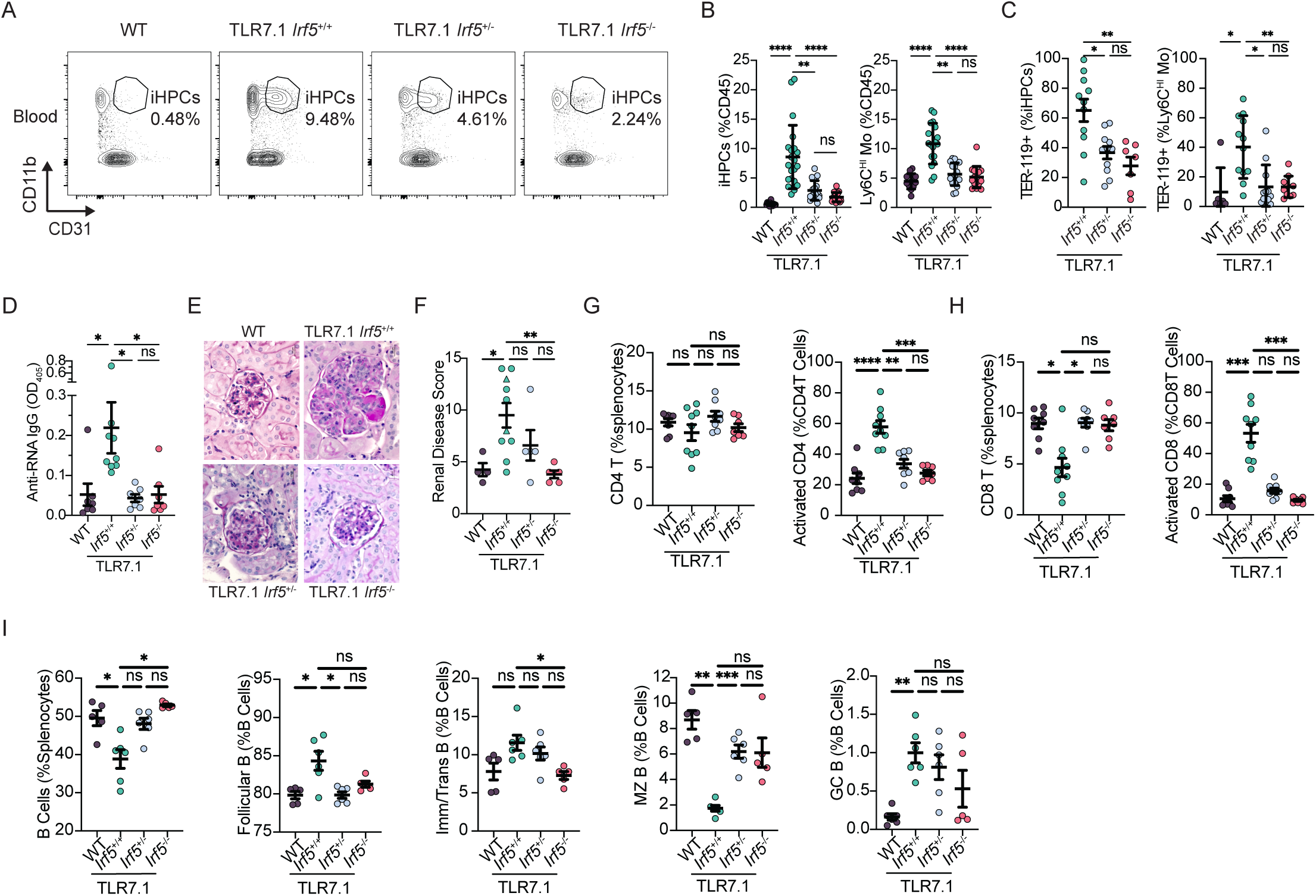
In vivo, IRF5 is required for iHPC differentiation and function, and its loss disrupts B and T cell compartments. (A-I) MAS-like and lupus-like disease and immune cell populations were assessed by flow cytometry in the blood and spleens of WT, TLR7.1 *Irf5*^+/+^, TLR7.1 *Irf5*^+/−^, and TLR7.1 *Irf5*^−/−^ mice flow cytometry at 14-16 weeks of age (n=8-10 per group). (A) Representative gating of iHPCs as a percent of CD45^+^ cells in the blood after gating on live CD45^+^Ly6G^−^SSC-A^LO^ cells (full gating shown in Fig. S1 C). (B) iHPCs and Ly6C^HI^ monocytes as percent of CD45^+^ cells in blood at 14-16 weeks of age. (C) Percentage of iHPCs and Ly6C^HI^ monocytes with intracellular RBCs in blood at 14-16 weeks of age. (D) Serum anti-RNA antibodies were measured by ELISA from mice at 14-16 weeks of age (n=7-9 per group). (E-F) Kidney pathology at 36 weeks (circles) or at the terminal point (triangles) was assessed by PAS staining. (E) Representative images and (F) renal disease score. (G) Splenic CD4 T cells as a percentage of total splenocytes; percentage of CD4 T cells that were CD44^+^CD62L^−^ activated/effector cells. (H) Splenic CD8 T cells as a percentage of total splenocytes; percentage of CD8 T cells that were CD44^+^CD62L^−^activated/effector cells. (I) Splenic B cells as a percentage of total splenocytes; Follicular, Immature and Transitional, Marginal Zone, and Germinal Center B cells as a percentage of total splenic CD19^+^ B cells. *p< 0.05, **p<0.01, ***p<0.001, ****p<0.0001. Kruskal-Wallis test (B-D, F, G (Total CD4 T cells), H (Total CD8 T cells), I (Fol. and Imm/Trans)); Brown-Forsythe and Welch ANOVA test G (activated CD4 T), H (activated CD8 T); I); each symbol represents an individual mouse; mean +/− SEM shown.

**Fig. S5:**
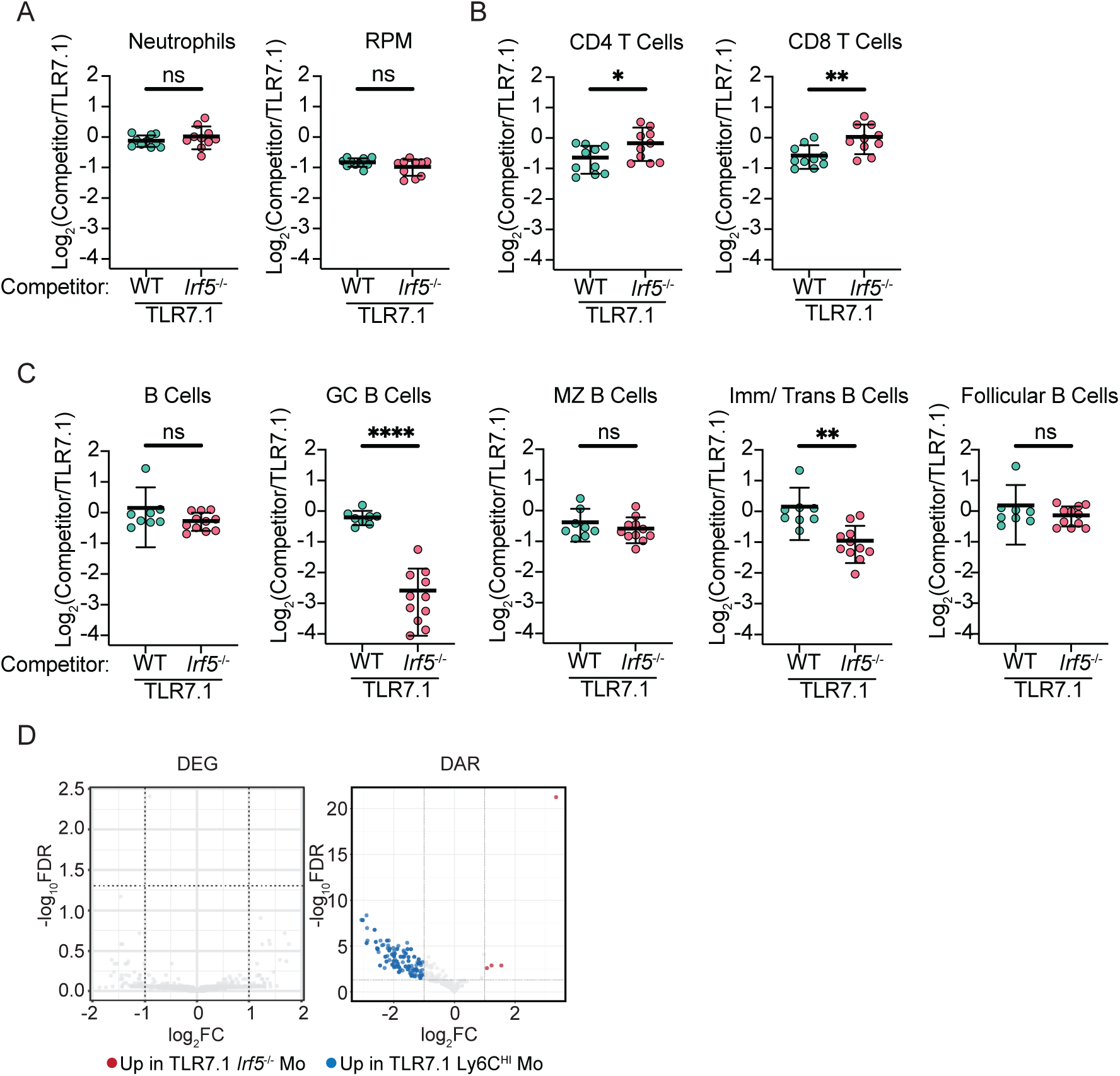
Cell-intrinsic IRF5 is required for iHPC differentiation. (A-C) Ratio of CD45.2 (TLR7.1 *Irf5*^−/−^ or TLR7.1) to CD45.1.2 (TLR7.1) cells in spleens from the chimeras described in Fig. 4. (A) Neutrophils and red pulp macrophages (RPM), (B) CD4 and CD8 T cells, (C) Total CD19^+^ B cells, GC B cells, MZ B cells, Imm/Trans B cells, and Fo B cells. (D) Paired bulk-RNA seq and ATAC seq of TLR7.1 Ly6C^HI^ monocytes (n=10-17) and TLR7.1 *Irf5*^−/−^ Ly6C^HI^ monocytes (n=5-10) sorted from TLR7.1:TLR7.1, and TLR7.1:TLR7.1 *Irf5*^−/−^ chimeras. Volcano plots showing the DEGs and DARs in TLR7.1 Ly6C^HI^ monocytes vs. TLR7.1 *Irf5*^−/−^ Ly6C^HI^ monocytes (log_2_Fc ≥ 1, adj.p <0.05 (blue up in TLR7.1 Ly6C^HI^ monocytes, red up in TLR7.1 *Irf5*^−/−^ Ly6C^HI^ monocytes). ns, p>0.05, *p<0.05, **p<0.01, **p<0.0001, Student’s unpaired t-test (A-C). Each symbol is an individual mouse, mean +/− SEM shown.

**Fig. S6:**
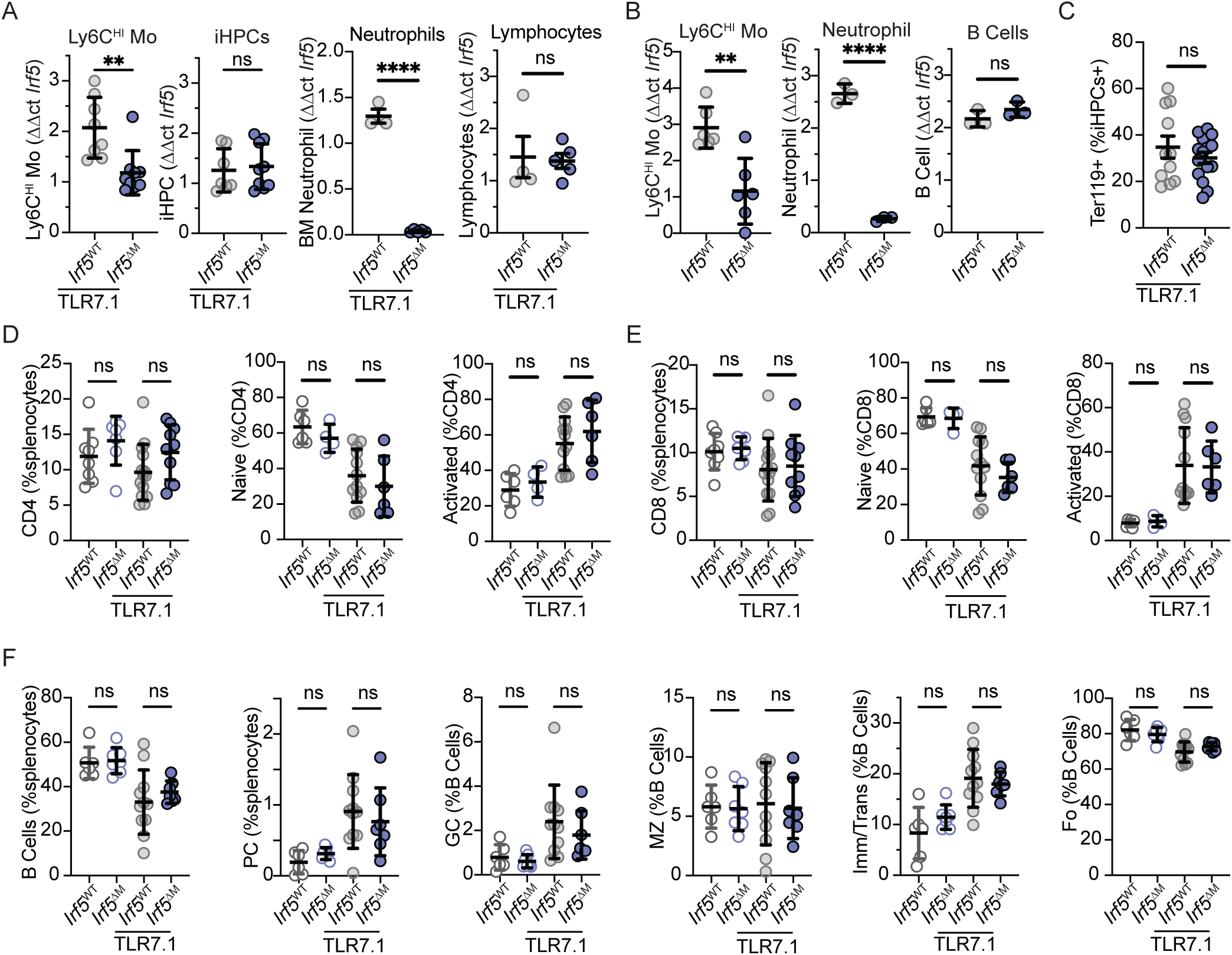
Myeloid cell *Irf5* deletion is dispensable for B and T cell alterations in TLR7.1 mice; cell-intrinsic IRF5 in splenic myeloid and lymphoid populations. (A, C-F) Analysis of TLR7.1 *Irf5*^ΔM^ and TLR7.1 *Irf5*^WT^ mice or (B) *Irf5*^ΔM^ and *Irf5*^WT^ mice. (A) *Irf5* expression by qPCR in sorted from splenic iHPC, splenic Ly6C^HI^ monocytes, BM neutrophils, and splenic lymphocytes from TLR7.1 *Irf5*^ΔM^ and TLR7.1 *Irf5*^WT^ mice (n=3-5 per group). (B) *Irf5* expression by qPCR in sorted splenic Ly6C^HI^ monocytes, neutrophils, and lymphocytes from *Irf5*^ΔM^ and *Irf5*^WT^ mice (n=3-6 per group). (C) iHPC RBC phagocytosis measured as percent iHPC with intracellular RBCs by flow cytometry at 14-18 weeks of age (n=12-15 per group). (D-F) Splenic T and B cells by flow cytometry at 14-18 weeks of age (n=8-15 per group). (D) Splenic CD4 T as a percentage of total splenocytes and CD44^−^CD62L^+^ naïve and CD44^+^CD62L^−^ activated cells as a percentage of CD4 T cells. (E) Splenic CD8 T as a percentage of total splenocytes and CD44^−^CD62L^+^ naïve and CD44^+^CD62L^−^ activated cells as a percentage of CD8 T cells. (F) Splenic CD19^+^ B cells and CD138^+^ plasma cells as a percentage of total splenocytes. Germinal center (GC) B cells, marginal zone (MZ) B cells, immature and transitional (Imm/Trans) B cells, and Follicular (Fo) B cells as a percentage of total splenic CD19^+^ B cells. **p<0.01, ****p<0.0001, Welch’s t-test (A-C); Brown-Forsythe and Welch ANOVA test (D, E (Total CD8 T cells), F (Total, MZ, Imm/Trans, Fo B Cells); Kruskal-Wallis test (E, F); Each symbol is an individual mouse, mean +/− SEM shown.

**Fig. S7:**
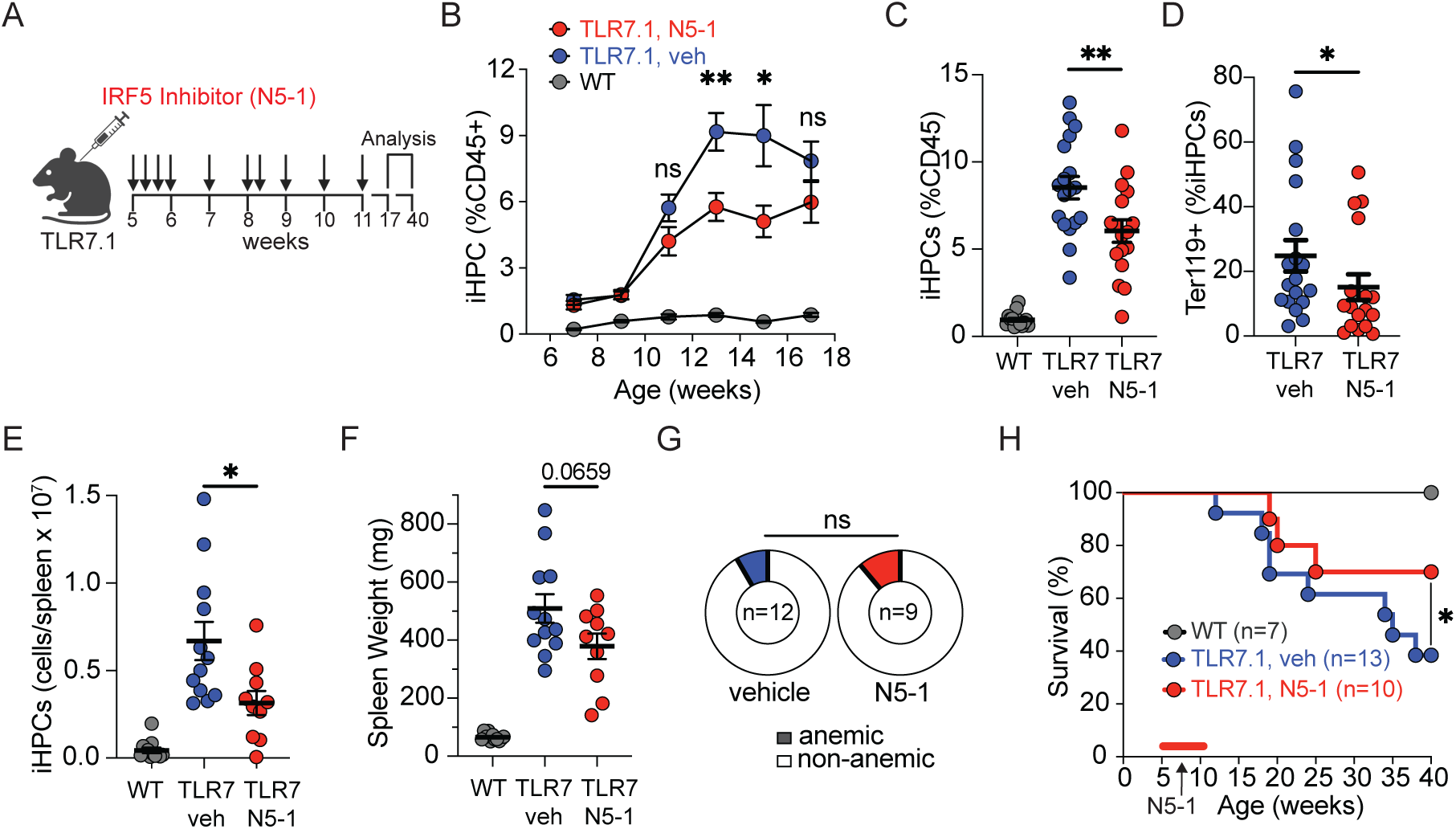
Preventative treatment of TLR7.1 mice with the IRF5 inhibitor N5-1 reduces iHPCs and MAS. Mice were treated 10 times with N5-1 or vehicle (PBS) between 5 and 11 weeks of age and assessed for blood iHPCs and MAS-like disease. (A) Schematic of preclinical N5-1 treatment of TLR7.1 mice. (B-D) iHPCs and iHPC RBC phagocytosis were assessed in the blood every two weeks from 7 to 17 weeks of age (WT n=6, N5-1-treated TLR7.1 n=11, vehicle-treated TLR7.1 n=11). (B) Blood iHPCs over the course of 7 to 17 weeks. (C-F) Blood and splenic iHPCs and MAS-like disease was assessed in a cohort of N5-1 or vehicle treated TLR7.1 mice at 17 weeks of age. (C-D) Blood iHPCs as a percent of CD45^+^ (C) and percent Ter119^+^ iHPCs (D) (N5-1 TLR7.1 n=9, vehicle-treated TLR7.1 n=12, WT n=7). (E) Splenic iHPCs and (F) spleen weights (WT n=12, N5-1 TLR7.1 n=10, vehicle-treated TLR7.1 n=12). (G) Percentage of mice with moderate-severe anemia (RBC <7 M/μl) in pre-clinical N5-1 treated and vehicle-treated TLR7.1 mice by 17 weeks of age (N5-1 TLR7.1 n=9, vehicle-treated TLR7.1 n=12). (H) Kaplan-Meier survival curve showing survival of WT (n=7), preclinical N5-1 treated TLR7.1 (n=10), and control vehicle treated TLR7.1 mice (n=13) over 40 weeks. *p<0.05, **p<0.01, Welch’s t-test (B-D); Mann-Whitney test (E-F); Wilson-Brown Binomial test (G). Log-rank (Mantel-Cox) test (H). Each symbol is an individual mouse.

**Table S1.**
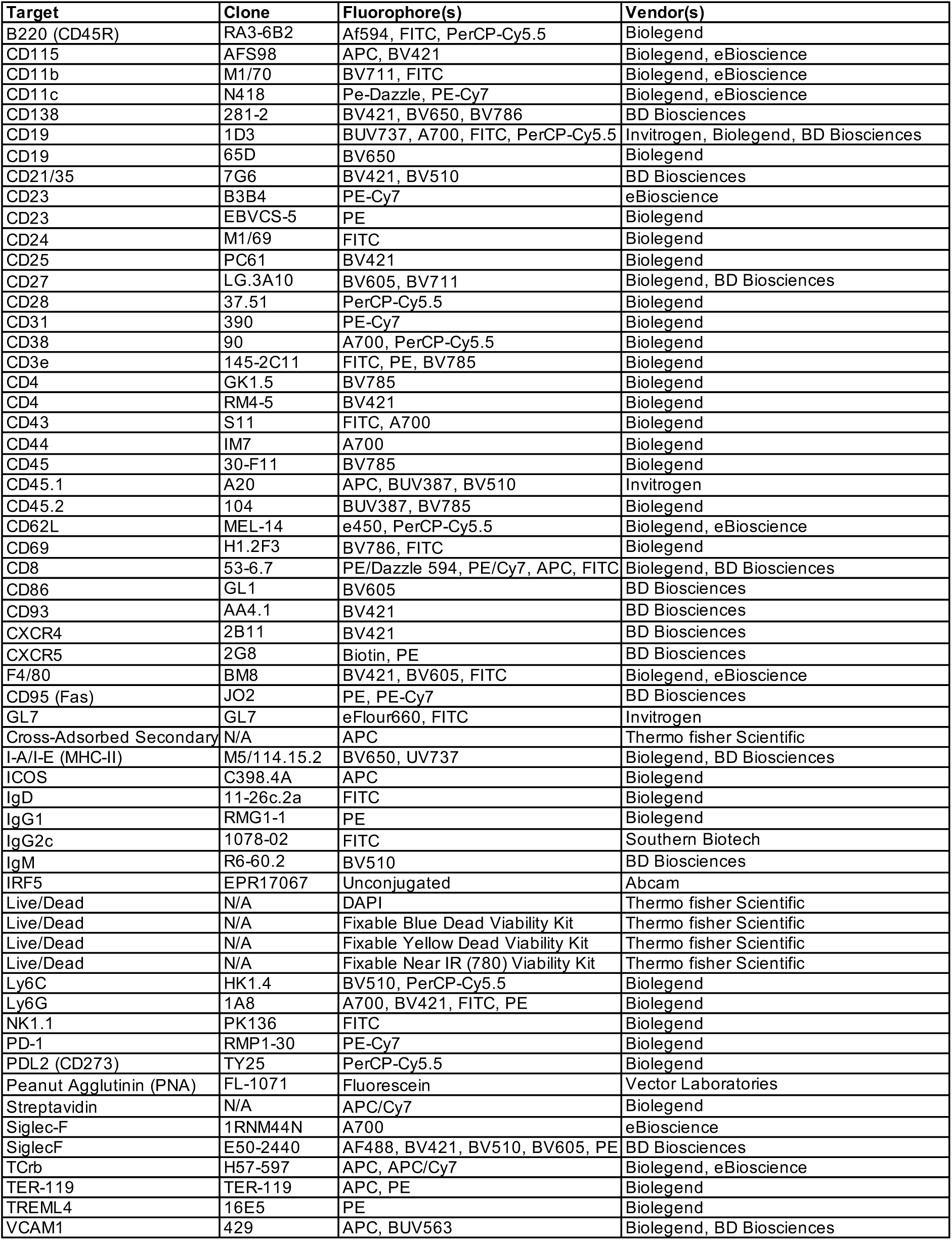

